# Prolonged low flows and non-native fish operate additively to alter insect emergence in mountain streams

**DOI:** 10.1101/2024.11.03.621706

**Authors:** Charlotte Evangelista, Mathieu Buoro, Kyle Leathers, Tatiana Tronel, Stephanie M. Carlson, Albert Ruhi

## Abstract

Climate-induced flow alteration is subjecting mountain streams to more frequent and severe low-flow periods due to lower snowpack and earlier snowmelt. Yet, anticipating how stream ecosystems respond to prolonged low flows remains challenging because trophic levels can respond differently, and non-native predators could dampen or amplify responses. Here, we conducted a large-scale experiment to examine how early, prolonged low flows projected by the end of the century in California’s Sierra Nevada will alter mountain stream food webs and emerging insect flux—a critical stream-to-land cross-ecosystem linkage. Additionally, we tested whether Brown trout (*Salmo trutta*), a widespread non-native top predator, would change food-web responses to low-flow conditions. We found that early low flows and non-native fish effects were additive rather than synergistic or antagonistic. Early low flows did not alter the overall rate of emerging insects but they did shift community structure and reduce the prevalence of small-sized individuals—possibly reflecting larger size at emergence and faster growth rates due to warming. In contrast, non-native fish presence increased seasonally-aggregated abundance of stream insects up to 12%, mainly by increasing abundance of Chironomidae and small-sized Ephemeroptera and Trichoptera. In channels with fish, benthic algal biomass doubled and scraper-grazer and collector-gatherer insects emerged 60% and 55% more than channels without fish, likely benefiting from trout keeping mesopredators at bay. This experiment illustrates that prolonged low flows and invasions can profoundly alter mountain river food webs even when operating additively; and shows how mesocosm-based research may help understand global-change driven disruption of cross-ecosystem linkages.

## INTRODUCTION

Climate change is altering freshwater ecosystems globally by shifting hydrologic cycles, snow and ice dynamics, and biota–via local extirpations, invasions, and shifts in organismal ranges (Rolls et al. 2012). In mountain ranges globally, anthropogenic climate warming has already reduced snowpack and advanced snowmelt, resulting in altered stream hydrographs characterised by earlier flow recession to baseflow levels (Parmesan et al. 2022). Seasonal and supra-seasonal droughts are expected to increase drastically in both length and severity over the next few decades regardless of the greenhouse gas emissions scenario (Siirila-Woodburn et al. 2021). In fact, the timing of mid-point runoff (i.e., the day when half of the year’s runoff has already occurred) projected under end-of-century climate change scenarios is anticipated to shift by over a month (and up to 6) in mountain ranges globally (Döll and Zhang 2010, Muelchi et al. 2021, Wieder et al. 2022). While mountain streams are expected to experience more frequent and severe low flow conditions (defined here as baseflow conditions over the dry season, Smakhtin 2001), we do not yet understand how these conditions will impact stream communities–and the ecosystem processes that these communities control.

Prolonged low flows can impact stream food webs both directly and indirectly (Palmer and Ruhi 2019). Stream invertebrate communities exposed to drought often change in taxonomic composition, with drought-sensitive taxa being replaced by drought-tolerant taxa (Ledger et al. 2013a, Herbst et al. 2019). Even when community composition does not drastically change, flow regimes can exert pressure on life history, morphological, and behavioral traits of aquatic ectotherms (Lytle and Poff 2004). For instance, most aquatic insects are biphasic and emerge as adult flying stages, and warming can induce earlier emergence at smaller sizes (Hogg and Williams 1996). However, some studies reported a lack of change in emergence timing (Brown et al. 2012), while others have shown earlier emergence at larger sizes in warmer sites (Gregory et al. 2000), or even taxon-specific responses to warming (Li et al. 2011). The variety of response types observed demonstrates the complexity of predicting how warming influences growth and development (Leathers et al. 2024). Hydrologically-driven changes in the composition and phenology of invertebrate communities can reshuffle trophic interactions; this can affect both the higher trophic levels via bottom-up processes (McIntosh 2022) and the base of the food web, via top-down control and associated trophic cascades (Power et al. 2008). For instance, warming due to prolonged low flows tends to enhance benthic algal production available to grazing invertebrates (Palmer and Ruhi 2019), with severity and timing of the dry and wet phases influencing basal resource responses (e.g., Truchy et al. 2020). However, because stress may differentially affect predators (Mor et al. 2022), changes in their abundance (e.g., their rarefaction) or performance (e.g., increased metabolic demands) can also propagate down the food chain via trophic cascades (e.g., by releasing or increasing pressure over primary consumers). Experimenting with complex communities composed of top predators and their prey may help advance our mechanistic understanding of how, and why, stream food webs respond to novel flow regimes (Walters and Post 2008).

In addition to hydrologic alteration, fish invasions have impacted headwater ecosystems in mountain regions globally (mainly salmonids; Crawford and Muir 2008). Many mountain headwater streams and lakes in Europe and the Western U.S. were once fishless, but from the mid-1800s, non-native salmonids were widely introduced to promote recreational fishing (Buoro et al. 2016). Introduced Salmonids can exert strong top-down control, leading to whole food-web change in terms of composition (Herbst et al. 2009, Buoro et al. 2016) and structure through size-selective predation (Blumenshine et al. 2000, Pope et al. 2009). Notably, their predation pressure can not only depress benthic invertebrate density but also greatly disrupt aquatic insect emergence–an important cross-ecosystem linkage between stream and riparian habitats (Baxter et al. 2004). However, prolonged low flows could affect the growth and performance of non-native salmonid predators and invertebrates–as well as interactions between the two. Specifically, higher metabolic rates of fish under warmer conditions can result in higher feeding rates (as long as the critical thermal maxima of fish is not exceeded, Cowles and Bogert 1944). Hence, it is critical to understand how emerging insect flux will respond to the combined effects of future low-flow regimes and non-native fish predation pressure–and to what extent responses may intensify as low-flow conditions advance.

Here, we conducted an outdoor mesocosm experiment in California’s Sierra Nevada to investigate how the flux of emerging aquatic insects may be altered by the individual and combined effects of prolonged low flows and non-native brown trout (*Salmo trutta*, hereafter referred to as *trout*). In California’s Sierra Nevada mountains, climate change-driven reductions in snowpack and earlier snowmelt are projected to advance mid-point runoff by up to six weeks by the end of the century under RCP8.5 (three weeks under RCP4.5; Reich et al. 2018). We simulated a “current” low-flow regime representative of contemporary conditions (hereafter, *current* scenario) and an end-of-century scenario where low-flow conditions start six weeks earlier (hereafter, *early* scenario treatment).Our experiment was designed to mimic mid- and high-elevation streams in California’s Sierra Nevada, which are naturally fishless (Herbst et al. 2009). Past experimental research mainly investigated community responses to flow magnitude (e.g., Ledger et al. 2013a), but the effects of low-flow timing and duration remain uncertain (but see Leathers et al., 2024). Here we built on these previous efforts by using a large, flow-through experimental channel system that allowed us to monitor complex food webs and a key cross-ecosystem linkage; while maintaining high replication levels (24 stream sections along 8 independent stream channels).

We expected that (*H1*) early low flow would increase the total abundance of emergent insect communities and alter their composition, especially by increasing the abundance of drought-tolerant taxa (Leathers et al. 2024). We also expected that (*H2*) top-down control exerted by introduced trout would alter emergent insect communities via selective predation on large-sized individuals and/or taxa (Blumenshine et al. 2000). Hence, we expected that (*H3a*) the combination of early low flow and presence of introduced trout would synergistically alter emergent insect communities (i.e., to a higher extent than the sum of individual impacts). Alternatively (*H3b*), predation effects could weaken in the prolonged low-flow treatment due to increasing abiotic stress on trout. Finally, we expected that (*H4*) the combination of early low flow and trout could trigger a trophic cascade, controlling grazing insects (primary consumers) and ultimately increasing benthic algae biomass. A robust test of these hypotheses should provide new insights into how climate-altered flow regimes affect food-web dynamics in vulnerable, high-mountain streams.

## MATERIALS AND METHODS

### Outdoor experimental channel setup

This study was conducted at an outdoor stream mesocosm facility located at the Sierra Nevada Aquatic Research Laboratory (SNARL, Mammoth Lakes, CA; 37° 37’ N, 118° 50’ W, elevation of 2200 m). The channels (50 m × 1 m) consist of alternating sections of riffle and pool habitat (6 pools and 7 riffles, Fig. 1A). The channels are made of concrete sealed with epoxy resin and the streambeds consist of a 0.3 m thick layer of silt, sand, gravel, and pebbles. This flow-through system is permanently sourced by water derived from Convict Creek, an adjacent perennial oligotrophic headwater stream that flows from the snowmelt-fed Convict Lake. Water supply can be independently controlled for each channel through a sluice gate located at the head of the channel (Fig. S1). These features maximise physical and ecological realism, which combined with high replication, have facilitated a suite of studies exploring aspects of invertebrate and fish ecology in the past (e.g., Jenkins et al. 1999, Saffarinia et al. 2022, Leathers et al. 2024). In our case, natural colonisation by organisms occurred for over three years prior to the start of the experiment. Invertebrate drift from the natural stream and feeder channel was not considered because past research showed drift rates in this system to be negligible over short timescales (Saffarinia et al. 2022).

**Figure 1.**
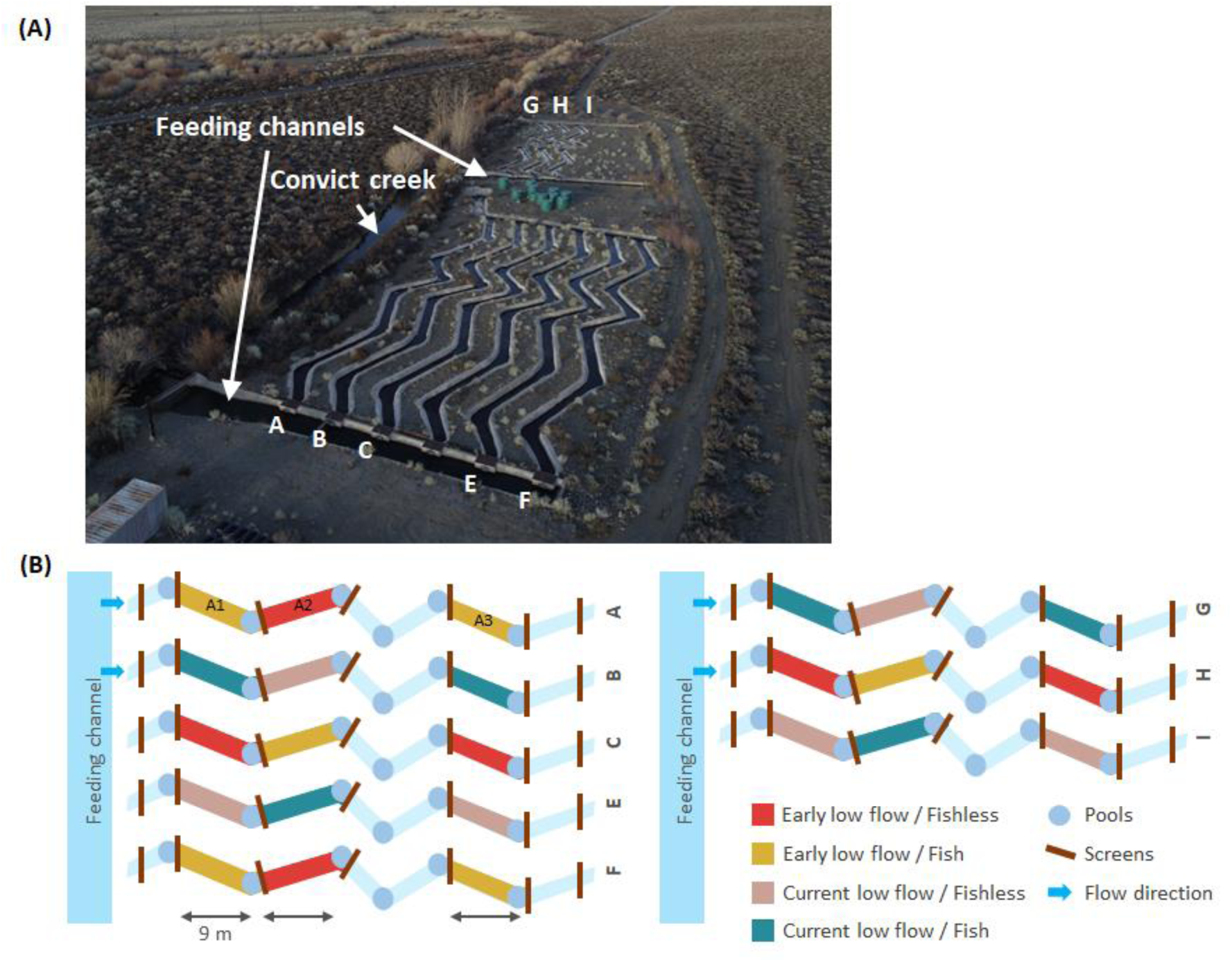
**(A)** Aerial view of the eight experimental channels (noted from A to I). Photographs: Paul Page. **(B)** Schematic presentation of the experimental setup with the four treatment combinations and their position within the experimental channels.

We used eight experimental channels arranged in two blocks, and each channel was divided into three subsections, for a total of 24 experimental units (Fig. 1). Each experimental unit was delimited by wire mesh screens (mesh size = 12.5 mm) that prevented trout from leaving the experimental units. All experimental units were 9 × 1 m and were composed of one riffle (upstream) and one pool (downstream; Fig. S1.A). We ensured trout had adequate habitat complexity and shelters in each experimental unit by adding large substrates (i.e., four boulders in each riffle and two cinder blocks along with boulders in each pool; Fig. S1). Prior to the experiment, successive electrofishing passes were carried out to capture pre-existing fish from the channels and release them back to Convict Creek. Finally, wire mesh screens were placed at the upstream and downstream ends of each channel to exclude any new fish from entering the experimental stream channels during the course of the experiment.

### Experimental design

The experiment consisted of a 2 × 2 factorial design with low-flow duration (*early* vs. *current*) crossed with presence of non-native trout (*fish* vs. *fishless*). Each of the four treatment combinations was replicated six times (Fig. 1B). The experiment comprised most of the growing season, starting on May 4th, 2022, when trout were introduced in the experimental units (see details below), and running for three months until August 2nd, 2022. Each channel was assigned to one low-flow treatment (i.e., *current* or *early*), then within each channel the three experimental units were assigned to the trout presence (*fish*) or absence (*fishless*) in a randomised block design (Fig. 1B). Because both factors should alter benthic invertebrate development (e.g., insect larvae and pupae; Blumenshine et al. 2000, Woodward et al. 2012), we anticipate these effects likely propagate to the emergence of flying adults (Statzner and Resh 1993).

We simulated realistic hydrographs for the *current* and *early* low-flow treatments. The *current* low-flow treatment was based on the contemporary hydrograph at Convict Creek (using US Geological Survey gage 10265200), with the flow regime reaching baseflow conditions in late July. The *early* low-flow treatment mimicked projected hydrologic conditions in Sierra Nevada’s streams under an end-of-century RCP8.5 scenario, with the flow regime reaching baseflow conditions six weeks earlier than in the current low flow treatment (Reich et al. 2018). Flow was gradually reduced over a week, and flow reduction was completed by June 6th, 2022 or by July 18th, 2022 in channels experiencing *early* or *current* low-flow conditions, respectively (Fig. 2A). Therefore, the experiment can be divided into three phases: *Acclimation phase*, when trout were introduced and flow levels increased uniformly in all channels (from May 4th to June 5th); *Phase I*, when flow was reduced in the *early* low-flow channels (from June 6th to July 17h); and *Phase II*, when flow was reduced in the remaining four channels (from July 18th to August 2nd; Fig. 2A). During *Phase I*, discharge (L s^-1^) differed by one order of magnitude between the *early* and *current* low-flow treatments (F_6,316_ = 207.2, P < 0.001; mean ± SE: *early* low flow = 5.4 ± 1.09, *current* low flow = 46.0 ± 7.2; Fig. 2). Discharge increased similarly and remained comparable between low-flow treatments during *Acclimation* and *Phase II*, respectively (*Acclimation*: F_6, 232_ = 2.29, P = 0.181, *early* low flow = 27.8 ± 6.2, *current* low flow = 34.10 ± 5.7; *Phase II*: F_6, 101_ = 3.6, P = 0.105, *early* low flow = 4.6 ± 0.5, *current* low flow = 7.3 ± 1.3; Fig. 2A). Streambed substrate remained submerged throughout the experiment, even during low-flow conditions (Fig. 2B).

**Figure 2.**
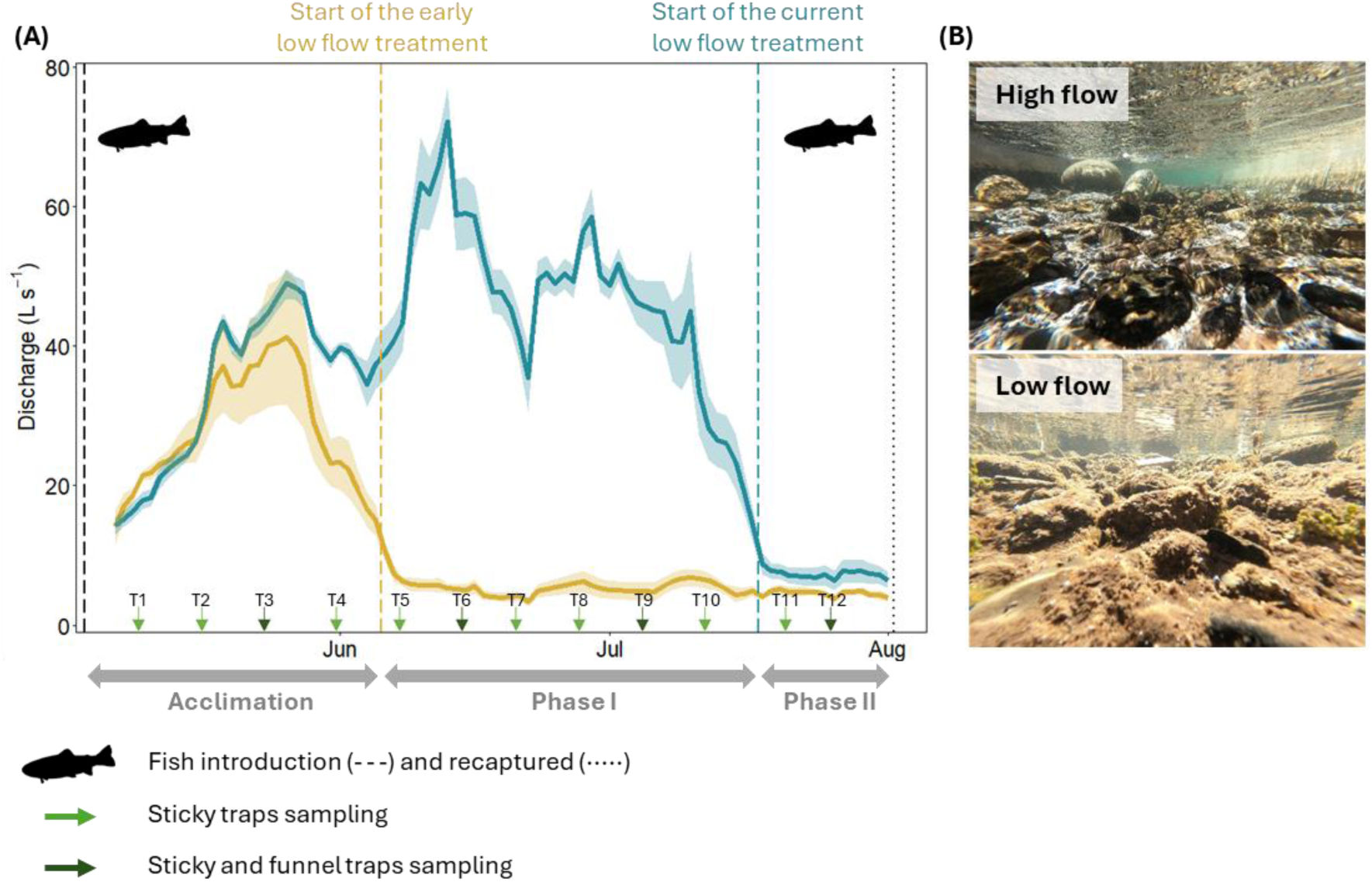
**(A)** Timeline of the experiment starting May 4 when fish were introduced and ending August 2. The experiment comprised three phases: an *Acclimation phase* where the predation treatment was applied and discharge increased similarly in all channels (from May 4 to June 5), a *Phase I* when discharge was reduced in the *early* low-flow treatment while maintaining constant in the *current* low-flow treatment (from June 6 to July 17), and a *Phase II* when discharge was reduced in the *current* low-flow treatment (from July 18 to August 2). Daily mean discharge averaged across channels receiving *current* vs. *early* treatment (continuous lines), and associated standard errors (shaded areas) are represented. Throughout the experiment, emerging insect communities were collected weekly using sticky traps (light green arrows) or combination of sticky and funnel traps (dark green arrows). **(B)** Underwater vision of a riffle during high and low-flow conditions.

Finally, half of our experimental units received Brown trout, a salmonid fish that was introduced to the region in the 1890s and has a large population in the Convict Creek watershed (Jenkins et al. 1999). Brown trout included in the experiment were captured via electrofishing from Convict Creek on May 4th, 2022. Individuals were then anaesthetized, measured for initial fork length (FL_i_ ± 1 mm), weighed (W_i_ ± 1 mg), marked with PIT-tags, and grouped into three size categories (small: 81-92 FL; medium: 130-140 FL; large:154-161 FL). On the same day, three trout from the three size categories were introduced into twelve experimental units (mean trout biomass = 24.00 ± 1.80 SD; trout density = 0.44 ind. m^-2^). These densities fell within the range of Brown trout density reported in the wild in the area (Jenkins et al. 1999).

### Monitoring of physical variables

Each channel was equipped with probes to measure water depth, temperature, and dissolved oxygen. All data were recorded every fifteen minutes throughout the experiment. In each channel, a HOBO U20L-04 pressure transducer (Onset Computer, Massachusetts, U.S.A.) was placed in the pool of the middle experimental unit. Two additional pressure transducers were placed on land to correct the data for fluctuations in atmospheric pressure, allowing for calculation of water depths. Depths were subsequently transformed into a time series of discharge values using channel-specific rating curves. Rating curves were developed for each channel by estimating discharge via channel depth measurements and velocity estimates taken with a Flo-Mate 2000 current meter (Marsh-McBirney Inc., Danaher Corp., Washington, D.C. U.S.A.). Rating curves were repeated 27 - 40 times per channel throughout the experiment. Water temperatures were measured in the pool microhabitat of each experimental unit with HOBO MX2202 sensors (Onset Computer, Massachusetts, U.S.A.). In turn, dissolved oxygen was measured in the middle experimental unit of each channel with a HOBO U26 probe (Onset Computer, Massachusetts, U.S.A.). All environmental data (i.e., discharge, temperature, and dissolved oxygen) were averaged to hourly values, which were then used to calculate daily means, minimums, and maximums per experimental unit (temperature) or per channel (dissolved oxygen). The diel range of temperature and dissolved oxygen were calculated by subtracting minimum from maximum values.

### Quantifying emergent insect flux, composition, and size structure

The emergent insect community was studied using sticky and funnel traps, which allowed for quantification of flux (individuals day^-1^) and for taxonomic identification of undamaged individuals, respectively. Funnel traps were floating devices consisting of pyramid-shape nets (177 µm mesh size), covering 0.33m^2^ of channel surface and equipped with a collector bottle (Fig. S1.D; Cadmus et al. 2016). Sticky traps were modified funnel traps with a collector bottle that was closed with tape, and with a piece of double-sided sticky tape (12.7 × 7.6 cm) attached to the inside of the pyramid frame (Fig. S1.D). For each experimental unit, one sticky trap was deployed weekly between the pool and the riffle to capture the influence of both riffle and pool habitats. After 24 hours of deployment, the sticky tapes were removed, photographed on both sides, and individually stored in a plastic bag. At the laboratory, glued emerging insects were counted and measured for total length (from tip of head to tip of abdomen) through photo analyses on ImageJ software. Count data were used to calculate both flux (individuals day^-1^) and cumulative flux of emerging insects, while body length data were used to calculate the insect community size spectra (*see details below*).

Because sticky traps are not well-suited for fine taxonomic resolution, funnel traps equipped with bottle collectors allowed for parallel collection of emerging insects that remained well preserved for taxonomic identification (Cadmus et al. 2016, Leathers et al. 2024). The funnel traps were deployed every three weeks (starting from May 24th; Fig. 2A), overlapping in space and microhabitat with the sticky traps. After 24 hours of deployment, emerging insects were gently collected with forceps, preserved in 70% ethanol, and later sorted to the family and subfamily level under a binocular microscope. Emerging insects collected using funnel traps were grouped into one of five functional feeding groups (i.e., scraper-grazers, collector-gatherers, collector-filterers, shredders, and predators).

Using the sticky trap data, we also characterised the community size spectrum, which describes the distribution of individual body sizes, in the emerging insect community (White et al. 2007). The size spectrum slope represents the relative proportion of small-sized versus large-sized individuals, with steeper slopes (i.e., more negative values) reflecting a decline of large-sized individuals, an increase of small-sized individuals, or both (Emmrich et al. 2011). Here, the size spectrum slope was fitted using individual body length of insects and the maximum-likelihood estimation method (MLE; Edwards et al. 2017). MLE performs better than traditional binning-based methods for calculating size spectra slopes when accounting for truncated power-law size frequency distributions and when sampling gear size selectivity is limited (Edwards et al. 2017), as is the case here. Specifically, we calculated community size spectra slopes of each treatment using the R package sizeSpectra (v.1.1.0, Edwards et al. 2020). Insects < 0.6 mm were very rare and were removed from the data set to avoid strong leverage. Because we were interested in the combined effects of predation by non-native trout and low-flow duration, the calculation did not include insects captured during the *Acclimation period* (i.e., when only predation occurred). To aid with interpretation of size spectra slopes, we visualised the length frequency distributions of emerging insects (Emmrich et al. 2011).

### Trout growth rates

Shortly after starting the experiment, we observed some trout movement beyond the experimental units. Consequently, on June 23rd we started electrofishing twice a week to confirm that the *fishless* experimental units and the ‘unused’ sections of the stream remained fishless. In addition, we confirmed trout presence in units with trout by placing an underwater camera (GoPro Hero 7) in the pool of each unit. These cameras were set up weekly for two hours before sunset, and footage was reviewed the day after.

On June 23rd, five trout were captured outside their experimental unit - in two unused units (one trout per unit) and in four *fishless* units (i.e., A2, B2, I1 and I3, one trout per unit). From June 27th to July 29th, five trout were captured in unused units of the channels, but *fishless* units always remained trout-free. All trout captured outside of their experimental units were released back to their original unit. This approach allowed us to maintain *fishless* (control) units and units with fish present. We also found four trout carcasses during the experiment, two of which (in unit A3) resulting from avian predation events that occurred during the last few days of the experiment, on July 24th and 28th. Dead trout were removed from the units but not replaced.

At the end of the experiment on August 2, trout were recaptured from our experimental units by electrofishing, checked for PIT-tags, weighed (W_f_ ± 1 mg), and then released alive back to Convict Creek. Recapture rate (n = 20; total recapture rate = 55.5%) declined with decreasing size of trout (83.3% for large-sized trout, 66.7% for medium-sized trout, and 16.7% for small-sized trout). In the experimental units subjected to the presence of non-native fish, 1 to 3 trout per unit were recaptured (mean = 1.75 ± 0.87 SD) in all units except for unit A3 where no trout were found at the end of the experiment (but note that two trout had died during the last week of the experiment in A3). One new trout (i.e., without a PIT-tag) was found in H2.

The specific growth rate (SGR, % month^-1^) of each recaptured trout was calculated as follows: SGR = 100 × (ln*W_f_* – ln*W_i_*) / Time

, where *W_f_* and *W_i_* are the final and initial weight, and Time is the duration of the experiment (3 months).

### Benthic algal biomass

At the end of the experiment (August 2nd), benthic algal biomass was measured *in situ* using chlorophyll-a concentration (µg cm^-2^) estimated using a portable fluorometric BenthoTorch probe that quantifies green algae, diatoms and cyanobacteria concentrations (BBE Moldaenke GmbH, Schwentinental, Germany). For each experimental unit, the probe was directly applied to the surface of 5 and 3 randomly chosen cobbles per riffle and pool habitat, respectively. These measurements were averaged to obtain unit-specific estimates of total benthic algae biomass per habitat.

### Statistical analysis

To characterise the abiotic conditions of the experimental streams, we tested the effects of prolonged low flow on daily values of temperature and dissolved oxygen (daily mean, daily minimum, daily maximum) using repeated measures ANOVA. We removed emerging insect data collected in units A2, B2, I1 and I3 before June 23th from subsequent analyses due to potential effects of trout movement.

We used linear mixed effects models (LMMs) to test the effect of treatments on emerging insect fluxes (abundance and cumulative abundance quantified using sticky traps) and on benthic algae biomass. We used generalised mixed-effect models (GLMMs) to test the effect of treatments on the most abundant taxonomic group (i.e., Order: Diptera, and Ephemeroptera-Trichoptera-Plecoptera pooled together, hereafter referred to as EPT; Family: *Chironomidae*, *Ceratopogonida*e, *Baetidae*, *Leptophlebida*e and *Hydroptilidae*) and functional feeding strategy (i.e., scraper-grazers, collector-gatherers, shredders, and predators), using count data from the funnel traps. We tested for treatment effects on community structure and composition via permutational multivariate ANOVA (PERMANOVA).

All models (LMMs, GLMMs and PERMANOVAs) included flow duration, predation and their interaction as fixed effects, as well as daily mean temperature (weekly integrated) as a covariate. LMMs and GLMMs also included *time* since the beginning of the experiment (i.e., the number of weeks) as a fixed effect. LMMs with emerging insects as response variables (sticky trap data) were tested over the three different phases (*Acclimation phase*: 4 sampling dates; *Phase I*: 6 sampling dates; *Phase II*: 2 sampling dates; one model per phase). In GLMMS, data collected from the *Phase I* and *Phase II* (3 sampling dates) were pooled together.

We performed all statistical analyses using R v.4.3.0 (R Development Core Team 2023). In particular, we fitted LMMs and GLMMs using the ‘lme4’ package (v.1.1.32, Bates et al. 2015) and the ‘glmmTMB’ package (v.1.1.8, Brooks et al. 2017), respectively. LMMs and GLMMs were fitted with stream identity as a random effect. Significance of fixed effects were tested using type III or type II Wald chi-square (χ^2^) or F tests performed with the ‘car’ package (v.3.1.2, Fox and Weisberg 2019). Significant interactions were further analysed using pairwise comparisons carried out using the ‘emmeans’ package (v.1.8.5, Length 2023). We found no evidence of collinearity among explanatory variables using the variance inflation factor (VIF). For LMMs, we also checked assumptions of linearity and homogeneity of variances on residuals using the ‘performance’ package (v.0.10.4, Lüdecke et al. 2021). To improve variance homogeneity and residuals normality we square-root and log transformed emerging insect abundance and algae biomass in LMMs, respectively. For each GLMM, the best distribution (i.e., negative binomial or generalised Poisson distributions) was chosen based on diagnostic plots performed with the DHARMa package (v.0.4.6, Hartig 2020).

PERMANOVAs were carried out in the ‘vegan’ package (v.2.6.4, Oksanen et al. 2022) by implementing Bray Curtis or Sørensen dissimilarities of the square root-transformed abundance of emerging insects. The zero-adjusted Bray Curtis dissimilarity was measured to enable calculations of empty samples by forcing a pair of blank samples to a dissimilarity of 0% (Clarke et al. 2006). Statistical tests performed using vegan’s ‘betadisper’ function indicated that one PERMANOVA deviated from homogeneity of multivariate dispersions (see details in the Results section). We used Non-metric multidimensional scaling (NMDS) plots to visualise effects of treatment combinations on community structure and composition. The ‘envfit’ function identified taxa that significantly contributed to community dissimilarity, then GLMMS were used to test the effect of treatment on the abundance of these particular taxa.

Hedges’s effect sizes were calculated to quantify the magnitude of the effects of prolonged low flows and the introduced fish on emerging insect metrics and benthic algae biomass. For each response, effects sizes were calculated as follows:

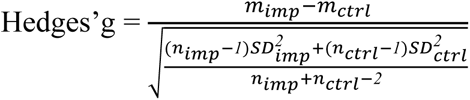

where *m* is the group mean and *SD* is the group standard deviation of the response variable determined as control (*ctrl*) and impact (*imp).* When calculating the effect sizes of prolonged low flows, data measured in the *current* low-flow conditions were used as control and data measured in the *early* low-flow conditions were used as impact. When calculating the effect sizes of introduced fish, data collected in *fishless* units were used as control, while those in units with non-native trout were used as impact. Following convention, the absolute mean effect size of each treatment was interpreted as negligible if |g| < 0.20, small if |g| < 0.30, medium if |g| < 0.80 and large if |g| ≥ 0.80 (Des Roches et al. 2019).

Finally, the effect of low-flow duration on specific growth and recapture rates of trout were tested using a linear model (LM) and a generalised linear model (GLM), respectively. LM included the initial size of individual trout as a covariate and GLM was fitted with a binomial distribution.

## RESULTS

### Abiotic and biotic environment

Prior to the experimental onset of low flows (Fig. 2), the different stream channels did not differ in their temperature and dissolved oxygen (DO) regimes (Table S1; Fig. S2). However, water temperature increased immediately with low-flow onset in *Phase I* (F_6,992_ = 18.19, P = 0.005), ultimately leading to an average 1.5°C increase in the daily range of water temperature in *early* relative to *current* treatments (Fig. S2.A). During *Phase I,* mean DO levels did not differ between treatments (F_6,320_ = 3.83, P = 0.098), but daily DO range (i.e., maximum – minimum values) was 1.2-fold higher in *early* than in *current* low-flow channels (F_6,320_ = 6.56, P = 0.043). This effect on DO variability was mainly driven by lower minimum DO values in the *early* relative to *current* low-flow channels (Fig. S2.B). During *Phase II* of the experiment, temperature, and DO values were similar between *early* and *current* low-flow channels (Table S1; Fig. S2). Overall, harmful hypoxia was not reached in the experimental channels (Fig. S2.B).

Data collected using funnel traps indicated that Diptera represented 88% of the total abundance of emerging insects, with the *Chironomidae* and *Ceratopogonidae* families composing 88% and 10% of the dipteran order. Within EPT, dominant families included *Baetidae* (35% of EPT), followed by *Hydroptilidae* (23%), and *Leptophlebidae* (19%). From a functional feeding group standpoint, communities were mainly dominated by collector-gatherers (67% of total abundance), followed by shredders (18%) and predators (10%). Scraper-grazers and collector-filterers represented only 4% and 1% of the total abundance of emerging insects, respectively

### Effects of prolonged low flows on the non-native predator

Trout recaptured at the end of the experiment (n = 20) had grown 41% faster in the *current* relative to *early* low-flow treatments (mean ± SE: 16.5 ± 2.5 and 9.8 ± 3.7, respectively), although this difference was not statistically significant (F_1,17_ = 2.72, P = 0.117; Hedges’g = −0.87 CI_95%_ [−1.84; 0.10]; Table S2, Fig. S3). Trout recapture rate was not influenced by the low flow treatment (F_1,10_ = 4.37, P = 0.063; Hedges’s effect size = −1.32 CI_95%_ [−2.71; −0.05]; Table S2).

### Effects of prolonged low flows and predation on emergent insect abundances

Emergent insect communities were similar across channels (LM: F_7, 72_ = 1.74, P = 0.113; PERMANOVA: F_7,12_ = 1.83, R^2^ = 0.56, P = 0.098) and treatments (Table S3) before the experimental onset of low flows, both in terms of abundance and structure.

There was no interaction between low-flow duration and presence of non-native trout on the abundance (including cumulative abundance) of emerging insects (Table S3; Fig. 3). However, we did observe an immediate, short-lived increase in the abundance of emerging insects with the onset of flow changes (Fig. S4.A). Additionally, the presence of non-native trout influenced the cumulative abundance (seasonally-aggregated) of emerging insects, but did so in an unexpected way – by increasing the cumulative abundance of emerging insects for *Phase I* (*fish*: 46.9 ± 3.5 SE, *fishless*: 44.6 ± 4.1 SE) and *Phase II* (*fish*: 169.4 ± 6.4 SE, *fishless*: 151.1 ± 9.7 SE), by 4% and 12% (respectively) in experimental units exposed to trout (*Phase I*: F_1,112.4_ = 19.00, P < 0.001; *Phase II*: F_1,36.5_ = 8.94, P = 0.005).

**Figure 3.**
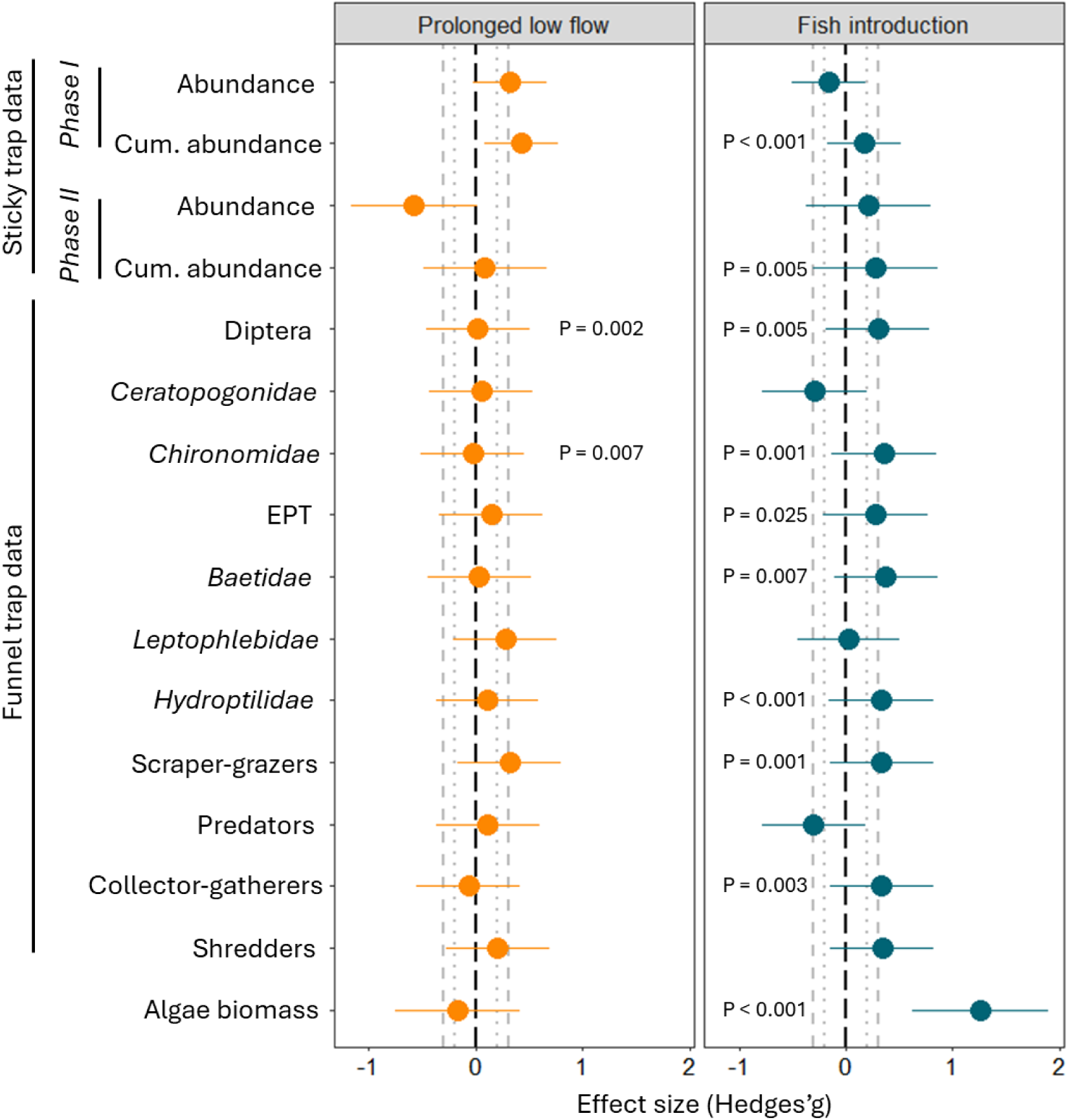
Effect size (Hedges’g) of prolonged low flows (left panel) and introduced fish (right panel) on the abundance of emerging insects and biomass of algae. Error bars represent 95% CI. Dotted and dashed grey lines represent a range of negligible (|g| < 0.2) and small (|g| < 0.3) effect size, respectively. P-values represent the results from mixed effects models (Table S3 and S4).

When looking at particular taxonomic groups (using funnel trap data), the cumulative abundance of Diptera was slightly higher in *early* than in *current* low-flow treatments (Hedges’g = 0.02; Table S4), but we detected no changes in EPT cumulative abundance (Table S4). Surprisingly, cumulative abundance of *Chironomidae* was 4% lower in *early* than in *current* low-flow treatments (mean ± SE; 24.3 ± 4.2 and 25.2 ± 5.7, respectively). In turn, the presence of non-native trout increased the cumulative abundance of both Diptera and EPT: Diptera were 39% higher in units with fish (Hedges’g = 0.30; mean_Diptera_ ± SE: *fish* = 33.3 ± 6.0, *fishless* = 24.0 ± 4.5), while EPT were 34% higher in units with fish (Hedges’g = 0.28; mean_EPT_ ± SE: *fish* = 5.9 ± 1.0, *fishless* = 4.4 ± 0.8) (Table S4; Fig. 3). The dipteran family that contributed the most to increasing cumulative abundance was *Chironomidae* (Hedges’g = 0.36; mean ± SE: fish = 29.7 ± 5.5, fishless = 19.5 ± 4.0), and variation in EPT abundance was mainly driven by increasing abundance of *Baetidae* (Hedges’g = 0.38; mean ± SE: fish = 2.1 ± 0.4, fishless = 1.3 ± 0.3) and *Hydroptilidae* (Hedges’g = 0.33; mean ± SE: fish = 1.4 ± 0.5, fishless = 0.6 ± 0.3) (Table S4; Fig. 3). Results using time-specific abundance of taxonomic and functional feeding groups mimicked results based on cumulative (seasonally-aggregated) abundance (Table S5).

When analysing responses by functional feeding groups (based on the funnel trap data), we observed that both the abundances and cumulative abundances of some feeding strategies were influenced by the non-native fish but not by the low-flow treatment (Table S4 and S5). For instance, the cumulative abundance of scraper-grazers (Hedges’g = 0.34; mean ± SE: fish = 2.1 ± 0.6, fishless = 1.1 ± 0.4) and collector-gatherers (Hedges’g = 0.33; mean ± SE: fish = 26.4 ± 4.8, fishless = 18.1 ± 3.6) was elevated under trout presence (Fig. 3), while predator abundance declined non-significantly (Hedges’g = −0.30; mean ± SE: fish = 2.5 ± 0.7, fishless = 4.0 ± 1.0) (Table S4; Fig. 3). Finally, the interaction between fish and low-flow treatment modulated the abundances of shredders (Table S4 and S5). Specifically, shredder cumulative abundance increased with the presence of non-native trout in the *early* low-flow conditions, but not under *current* low-flow conditions (Table S6; Fig. S5).

### Effects of prolonged low flows and predation on emergent insect composition

There was no significant interaction between low-flow duration and presence of non-native trout on emergent insect composition (Table S3). However, emerging insect community structure (Bray-Curtis distance) exhibited responses to low flows in *Phase I* (PERMANOVA: F = 4.71, P = 0.009; Fig. 4A). Specifically, the *Orthocladiinae* and *Prodiamesinae* subfamilies explained community dissimilarities between *current* and *early* treatments (Table S7). Similar responses were observed when considering community composition (Sørensen distance; F = 2.24, P = 0.022), but these results could be affected by non-homogeneous dispersion of the data (*betadisper*: P = 0.031). Indeed, communities from the *early* low-flow treatments appeared to be less variable (or dispersed) than those from the *current* low-flow treatments (Fig. 4B).

**Figure 4.**
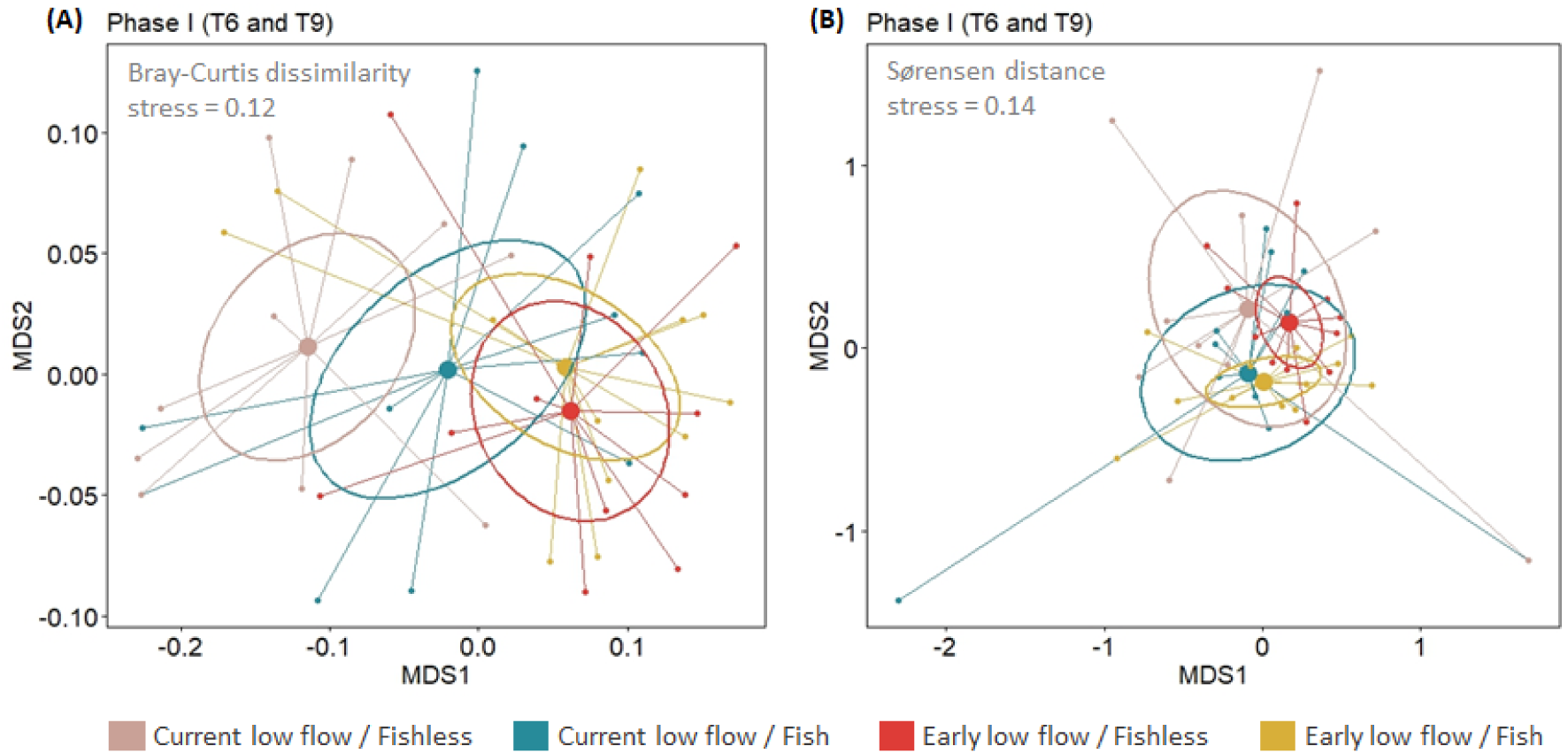
Nonmetric multidimensional scaling (NMDS) ordinations of emerging insect community **(A)** structure (Bray-Curtis dissimilarity; stress = 0.12) and, **(B)** composition (Sørensen distance; stress = 0.14) from the different treatment combinations. Samples (n = 48) were collected during *Phase I* of the experiment at T6 and T9. Large dots represent centroids for each treatment, whereas small dots represent insect communities within experimental units. Ellipses are based on 95% standard error around the treatment centroids.

During *Phase II*, the emerging insect community was similar among low-flow treatments (Table S3). Although previous analyses indicated that the cumulative abundance of some Families were influenced by the presence of trout (see above), multivariate analyses revealed no influence of non-native trout on the structure and composition of emerging insect communities (Table S3).

### Effects of prolonged low flows and predation on emergent insect size spectra

With regards to community size spectra slopes, we also did not observe a significant interaction between low-flow duration and presence of non-native trout. However, prolonged low flows flattened the slope of the emerging insect community from −1.31 (*early* low flow/fishless: CI_95%_ [−1.39; −1.23]) and −1.35 (*early* low flow/*fish*: CI_95%_ [−1.43; −1.27]) in the *early* low-flow treatments, to −1.57 (*current* low flow/*fishless*: CI_95%_ [−1.66; −1.49]) and −1.64 (*current* low flow/*fish*: CI_95%_ [−1.73; −1.56]) in the *current* low-flow treatments (Fig. 5A, Fig. S6). This result likely stems from a decrease in the prevalence of individuals < 1.5 mm in *early* compared to *current* low-flow treatments, associated with an increasing prevalence of individuals with body size ranging from 1.5 to 3.5 mm (Fig. 5B).

**Figure 5.**
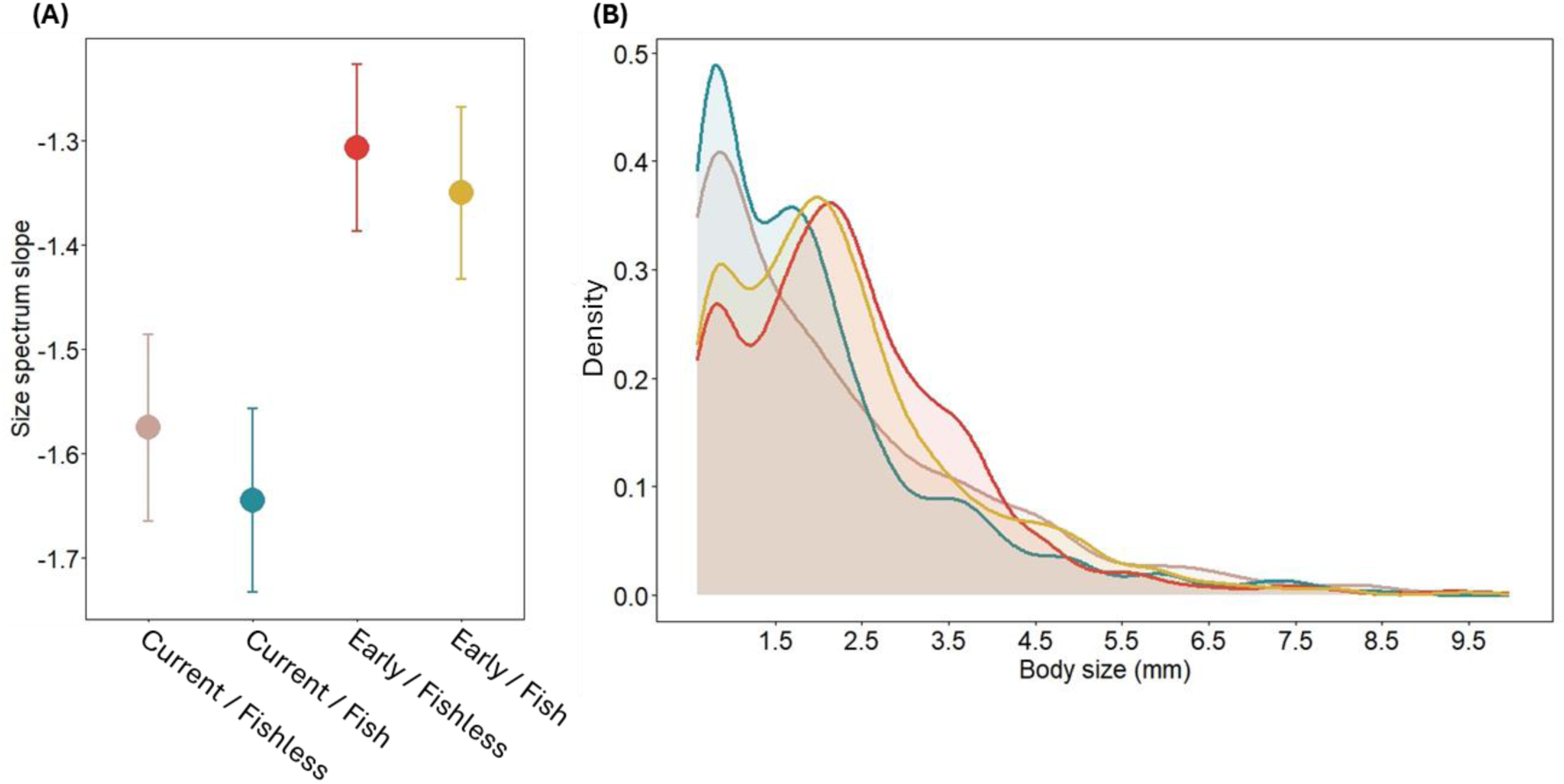
**(A)** Effect of treatment combinations on the size spectra slopes fitted using maximum likelihood estimation. Mean and 95% CI are represented. **(B)** Body size distribution of emerging insects in the different treatment combinations. Data collected during *Phase I* and *Phase II* are pooled together.

### Effects of prolonged low flows and predation on primary producer biomass

Finally, biomass of benthic algae was also not affected by the interaction of low-flow duration and presence of non-native trout (Table S3). However, although algal biomass was similar between *early* and *current* low-flow treatments (F_1,5.6_ = 0.91, P = 0.379; Fig. 6), trout presence did have strong effects (F_1,43.86_ = 24.13, P < 0.001), doubling benthic algal biomass in sections with trout relative to those without trout (*fish* = 0.96 ± 0.11 SE, *fishless* = 0.47 ± 0.02 SE; Table S3, Fig. 6). Changes in benthic algal biomass were mainly associated with an increased abundance of green algae and cyanobacteria (Table S3), which were 5- and 2-fold higher in the presence of non-native trout, respectively (mean_green algae_ ± SE: *fish* = 0.26 ± 0.09, *fishless* = 0.05 ± 0.01; mean_cyanobacteria_ ± SE: *fish* = 0.37 ± 0.07, *fishless* = 0.17 ± 0.03). However, abundance of epilithic diatoms barely responded to the fish treatment (mean_diatoms_ ± SE: *fish* = 0.34 ± 0.04, *fishless* = 0.25 ± 0.02).

**Figure 6.**
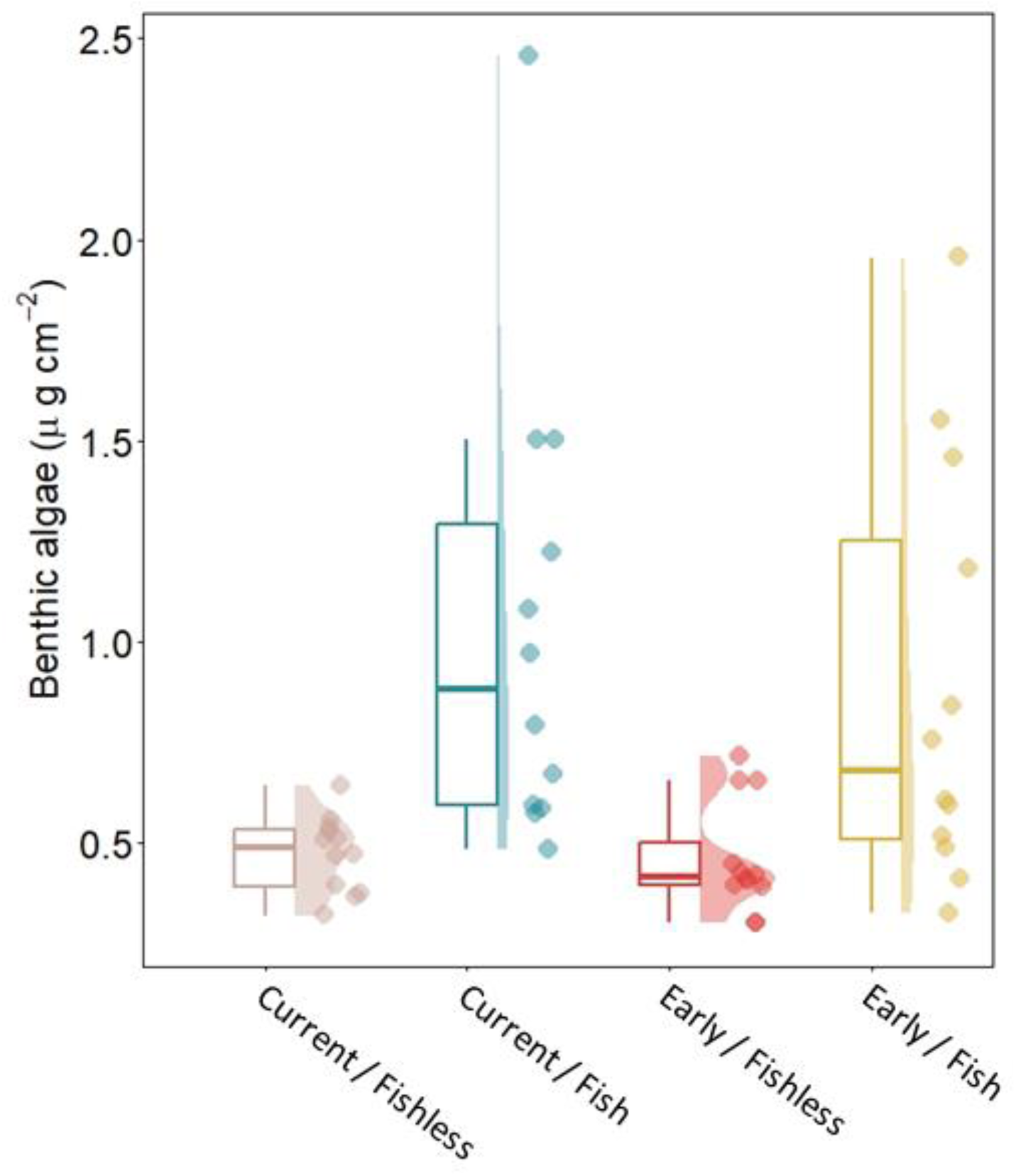
Raincloud plots showing the effect of treatment combinations on benthic algae biomass at the end of the experiment. Dots represent the samples (n = 12 per treatment), boxplots and half violin plots represent the probability density of the data.

## DISCUSSION

Several studies have experimentally shown the effects of climate-induced flow alteration on stream food webs (e.g., Ledger et al. 2013a, Ledger et al. 2013b, Aspin et al. 2019, Leathers et al. 2024). However, how mountain stream food webs are being impacted by the joint effects of climate change and non-native species, which are expanding their ranges 100 times faster than native species (Bradley et al. 2024), remains largely unknown. Here, we used an outdoor, experimental stream array to anticipate the joint effects of prolonged low flow scenarios and the presence of a global invader, the Brown trout. We focused on the riverine food web and emerging (adult) insects – a key linkage connecting streams to the adjacent terrestrial environment (Baxter et al. 2004). We found that prolonged low flows altered the structure of emerging insect communities and increased the abundance of some groups (Diptera) but not of the overall emerging assemblage (H1). Contrary to our second hypothesis (H2), we found that the abundance of the two major taxonomic groups (i.e., Diptera and EPT) increased with trout presence, mainly due to increased abundance of Chironomidae and small-sized Ephemeroptera (i.e., Baetidae) and Trichoptera (i.e., Hydroptilidae). Scraper-grazers and collector-gatherer insects also emerged more (an increase of 60% and 55%, respectively) in channels with fish, likely benefiting from mesopredators (e.g., *Doroneuria baumanni*, Turbellaria flatworms), presumably being kept at bay by fish predation (Herbst et al. 2009). However, the size structure of the emerging insect community (as measured by their spectra slopes) did not substantially change when exposed to fish. We also detected a reduced prevalence of small-sized insects in *early* low-flow channels, suggesting that small-medium sized individuals or species increased in abundance relative to the smallest individuals, and/or that insects emerged at larger size due to warming. We found that almost no response variable was influenced by the interaction between low-flow duration and the presence of non-native trout. This highlights that additive, rather than synergistic or antagonistic effects, governed responses (contrary to our third hypothesis, H3). Finally, benthic algae biomass did not differ between low-flow treatments but doubled in fish relative to the fishless treatment (H4). These findings stress that even when operating additively, climate change-induced low flows and invasion can strongly alter stream communities and disrupt important links to the terrestrial environment–with potential for responses to propagate beyond the riverine ecosystem (Leathers et al. 2024).

### Responses of emerging insects to climate-change induced low flows

Prolonged low flows were more strongly associated with changes in community structure than with changes in the overall flux of emerging insects. *Orthocladiinae* and *Prodiamesinae* subfamilies explained community dissimilarities in *current* versus *early* low flows. This result is consistent with the slight increase in the early low-flow channels in abundance of Diptera, a taxonomic group that is diverse in terms of thermal and dissolved oxygen tolerance, and includes species that are adapted to poor environmental conditions (Serra et al. 2016). This observation is consistent with previous studies (Leathers et al. 2024), but also masks taxon-specific responses. Indeed, contrary to Herbst et al. 2019, we found that *Chironomidae* (which represents 88% of all Diptera in our samples) were less abundant in *early* low-flow channels. We also did not detect any response of EPT to prolonged low flows, though the discharge in *early* low-flow channels during *Phase I* (mean = 5.4 L s^-1^ ± 1.09 SE) was within the range of values that caused EPT declines in natural headwater streams (Herbst et al. 2019). However, in our experiment, EPT communities were mainly composed of taxa with small body size (i.e. *Diphetor hageni*, *Baetis* sp., *Leptophlebidae,* and *Hydroptilidae*), which may be more resistant to drought due to lower metabolic demands and easier access to suitable habitat. Alternatively, low-flow conditions may not have been extreme enough to lead to stressful conditions, as illustrated by the fact that stream substrates remained wet throughout the experiment.

Ecological theory has long posited that smaller individuals are generally favoured under warmer conditions (“temperature-size rule”, Forster et al. 2012). Hence, one would expect size spectra slopes to be steeper after prolonged low flows due to warming-induced community shrinking (Yvon-Durocher et al. 2011). However, that was not the case here. Instead, the prevalence of medium-sized emerging insects (1.5– 3.5 mm) was higher in *early* compared to *current* low-flow conditions, resulting in flatter size spectra slopes. As reported previously, Chironomidae can become larger in terms of mean body size with rising temperature (Piggott et al. 2015). Therefore, the dominance of Chironomidae in our experiment could explain changes in size spectra slopes in response to low-flow duration, realized via temperature-dependent growth rates. Future experiments could reveal whether changes in size spectra slopes are caused by a shift in taxonomic composition or by individuals of the same species growing faster and to larger sizes. Taken together, these results confirm the importance of considering both size spectra metrics and body size distribution (e.g., size-frequency distribution) when evaluating effects of prolonged low flows on community size structure (Emmrich et al. 2011). Although our experiment was relatively short-lived (3 months), the main taxonomic groups were able to complete at least one generation. Investigating intergenerational or lagged effects of warming on community size structure is worthy of future research.

### Implications for cross-ecosystem linkages

Our results show that even if the low-flow treatment did not shift cumulative abundance of emerging insects, a short-lived response was apparent. In particular, the abundance of emerging insects increased immediately with the onset of low flows (Fig. S4.A). In the same study area, Leathers et al. (2024) found that during bird nesting, the presence of Brewer’s Blackbirds (*Euphagus cyanocephalus*) in the channels coincided with high production of emerging Chironomidae due to earlier decrease in discharge. Although those authors found a novel climate-driven prey-predator match, it is important to note that climate change often leads to phenological mismatches in trophic interactions (Thackeray et al. 2016). Phenological mismatches are particularly problematic in strongly-seasonal ecosystems such as high-mountain streams, where resources are transient. In addition, if earlier emergence is associated with changes in the nutritional composition of insects, it is likely that nutritional phenological mismatch will have harmful consequences for consumers (Twining et al. 2022). Therefore, further investigations are needed to track how climate change affects not only the biomass but also the nutritional quality of these cross-ecosystem subsidies (e.g., in terms of long-chain polyunsaturated fatty acids or LC-PUFAs).

### Effects of climate-change induced low flows on non-native trout

Cold-adapted salmonids often experience slower growth rates when exposed to drought and associated low flows (Harvey et al. 2006). In addition, long periods of low flows diminish habitat availability, making pools more suitable than riffles. For instance, Rosenfeld and Boss (2001) showed that during low-flow events, trout inhabiting riffles had lower growth rate than those inhabiting pools. In our experiment, trout growth decreased by 41% in *early* compared to the *current* low-flow treatments, even if limited sample size prevented us from detecting a statistically-significant effect.

Water turbulence can act as a shelter from visual hunting birds (Allouche and Gaudin 2001), suggesting that *early* low-flow conditions could have facilitated predation by shorebirds (e.g., herons). This could ultimately explain, at least partially, the lower recapture rate of trout in *early* low-flow conditions, which was particularly low when considering small-sized trout (16.7% compared to 83.3% and 66.7% for large- and medium-sized trout, respectively). Brown trout are territorial, and low-flow-induced habitat contraction likely boosted agonistic behaviour. We believe that small-sized trout, which were able to ‘sneak’ between pebbles, may have escaped from the fish-bearing experimental units to avoid harmful competition from large conspecifics.

### Top-down control in mountain stream food webs

Salmonids are known to influence fluxes of emergent stream invertebrates (e.g., Baxter et al. 2004). Here, the cumulative abundance of emerging insects was higher when trout were present, and this pattern was mainly driven by an increased abundance of *Chironomidae* midges and small-sized EPT (i.e., Baetidae mayflies and *Hydroptilidae caddisflies*). Top predators such as Salmonids can benefit small-sized insects if fish preferentially prey on invertebrate mesopredators (Pope et al. 2009). However, the slope of the size spectrum community was not influenced by the presence of trout, likely revealing that trout predation was not strongly size-selective. In contrast, we found that the effects of fish introduction differed among functional feeding groups, with substantial increases in collector-gathers, shredders, and scraper-grazers,, and slight declines in predators. While we cannot rule out the potential influence of repeated electrofishing of fishless units on invertebrate drift (Elliot and Bagenal 1972), changes in insect community composition and structure between fishless and trout units likely arose from trout keeping mesopredators at bay–thus releasing small insects from top-down control.

Consistent with previous studies, we found that algae biomass was greater in the presence versus absence of trout (e.g., Herbst et al. 2009). A trophic cascade could explain increased benthic algal biomass, but only if the increasing abundance of emerging scraper-grazer in the presence of trout was associated with a decreasing abundance of benthic scraper-grazer larvae. Hence, more information would be required to clearly identify whether changes in the composition of benthic (larval) communities reflect change in emergence of aquatic insects (Schmidt et al. 2013, Wesner 2013). An alternative explanation for the increased algal biomass is the role of fish, as large consumers, in nutrient recycling (Atkinson et al. 2017), which could ‘fertilize’ and enhance primary production. However, we believe this explanation is less plausible than the trophic cascade hypothesis, given the short residence time in the channels (approximately 2 and 11 minutes under high- and low-flow conditions, respectively; Leathers K., unpublished data).

### Additive effects of low-flow duration and non-native trout

Freshwater ecosystems are frequently impacted by multiple stressors, and interactive effects (synergistic or antagonistic) rather than additive effects often prevail (Jackson et al. 2016). For instance, Greig et al. (2002) found that stickleback predators (*Gasterosteus aculeatus*) strengthened earlier emergence of aquatic insects in warmer pond mesocosms, while Ross et al. (2022) found that the presence of sculpin (*Cottus nozawae*) buffered heatwave impacts on benthic macroinvertebrates. However, here we found no evidence for interactive effects between non-native fish and climate-induced prolonged low flows. Although frequently disturbed ecosystems are often more prone to invasions (‘environmental resistance hypothesis of invasion’, Moyle and Light 1996), the impacts of invaders on recipient ecosystems depend on many factors. Interactive effects could emerge between invaders and climate change in cases where climate change directly facilitates the establishment of the non-native species (‘alteration-invasion nexus’, Ruhi et al. 2019). In the studied area, brown trout were intentionally introduced in the 1890s to support recreational angling (Jenkins et al. 1999), meaning that (i) macroinvertebrates in the experimental streams were familiar with brown trout at some level, and (ii) the establishment of brown trout was independent of climate-induced hydrological alteration. Altogether, this could explain the absence of interactive effects in our stream experiment.

## Conclusions and future directions

Climate change and fish invasions are two major threats to mountain stream ecosystems globally. Yet, disentangling their effects remains challenging and often requires experimental approaches. Currently, most flumes and mesocosm systems have excluded large predators from food webs, because they cannot be housed in smaller experimental systems (Stewart et al. 2013). Large mesocosms include greater biological realism, but often lack adequate replication, which can ultimately limit inferences. It is also difficult to integrate downscaled climate change scenarios in stream experiments, and such mismatches can restrict realism and value of information gathered from experimental manipulations (Korell et al. 2019). Although any experimental approach has limitations, we believe our system balanced realism with control, providing mechanistic insight into how multiple stressors may alter mountain stream food webs in the future. Long-term experiments that capture intergenerational effects (e.g., Woodward et al. 2012), and more extreme flow scenarios (e.g., including flow intermittency and pool disconnection), could expand on and add context to our findings.

## ACKNOWLEDGMENTS

The work was performed at the University of California Valentine Eastern Sierra Reserves and we are grateful to Carol Blanchette, Jeremiah Eanes, and Ben Peck for logistical support. We also thank David Herbst for access to his lab; Stephane Glise for construction of floating traps; Parker Trimble, Amaïa Lamarins and Martin Dizel for field assistance; two anonymous reviewers for their suggestions that have improved the quality of the manuscript.

## ETHICS

The experiment was conducted under ACUC permit no. AUP-2021-11-14803 authorised by UC Berkeley and approved by UC Santa Barbara, and by California Department of Fish and Wildlife permit no. S-2122 30001-21319-001.

## DATA AVAILABILITY STATEMENT

The data and R code that support the findings of this study will be openly available in Figshare upon acceptance of the manuscript.

## AUTHOR CONTRIBUTION STATEMENT

**CE:** Conceptualization (equal); data curation (lead); formal analysis (lead); investigation (lead); methodology (equal); supervision (equal); visualization (lead); writing – original Draft Preparation (lead); Writing – review & editing (equal)**. MB:** Conceptualization (equal); funding acquisition (lead); investigation (equal); methodology (equal); project administration (equal); supervision (equal); writing – review & editing (equal). **KL:** Conceptualization (equal); investigation (supporting); methodology (equal); writing – review & editing (supporting). **TT:** Investigation (equal); methodology (supporting); writing – review & editing (supporting). **SMC:** Conceptualization (equal); funding acquisition (equal); investigation (supporting); methodology (equal); project administration (equal); writing – review & editing (equal). **AR:** Conceptualization (lead); funding acquisition (equal); investigation (supporting); methodology (equal); project administration (equal); supervision (equal); writing – original draft preparation (supporting); writing – review & editing (equal). All authors approved the final manuscript.

