## Supplementary material for "Prolonged low flows and non-native fish operate additively to alter insect emergence in mountain streams": Supplemetary material

**Supplementary figures and tables**

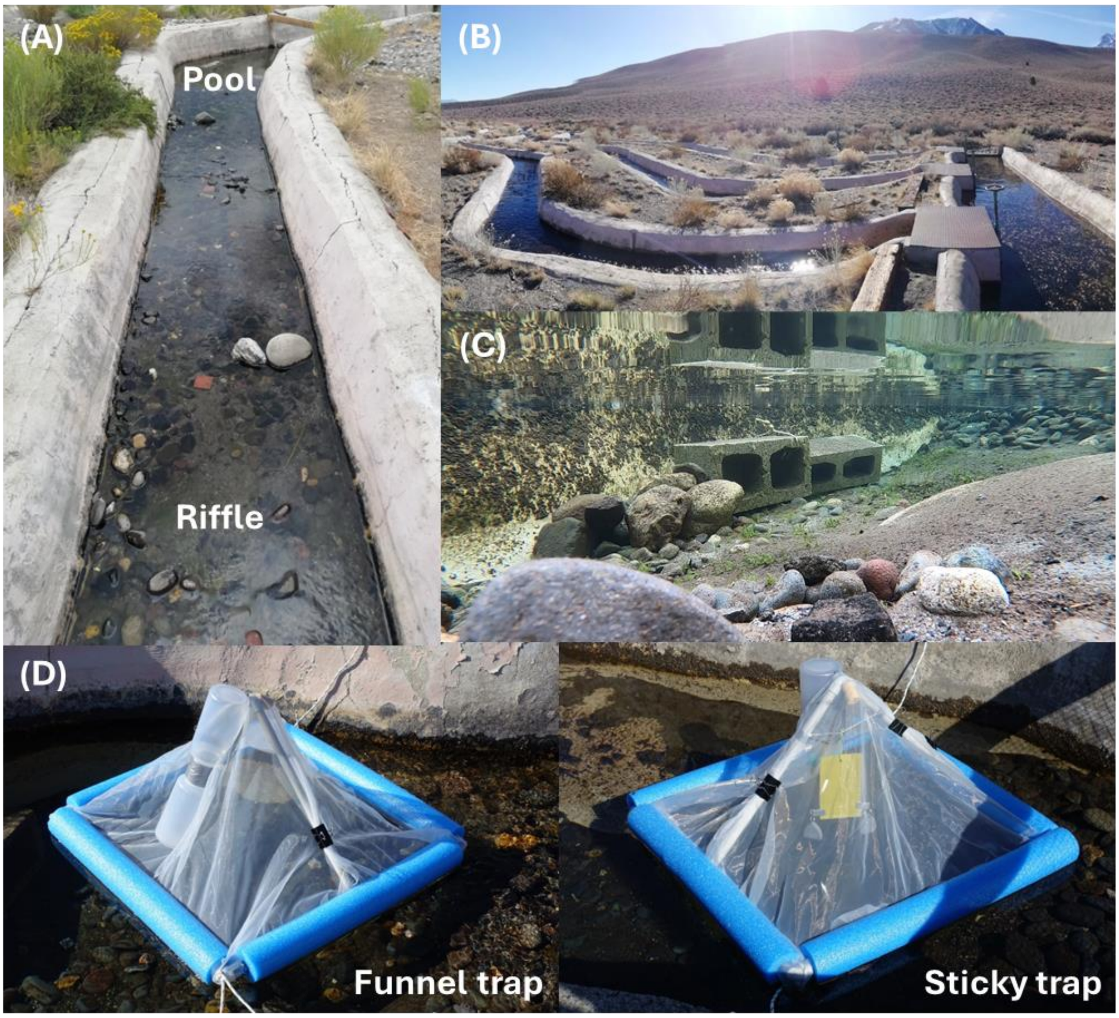

**Figure S1** Photographs of **(A)** an experimental unit composed by a pool and a riffle, **(B)** the head of the channels with sluice gates that allow to manipulate flow regimes, **(C)** underwater vision of a pool with shelters for fish and **(D)** the sticky and funnel traps used to collect emerging insects. Photographs: CE.

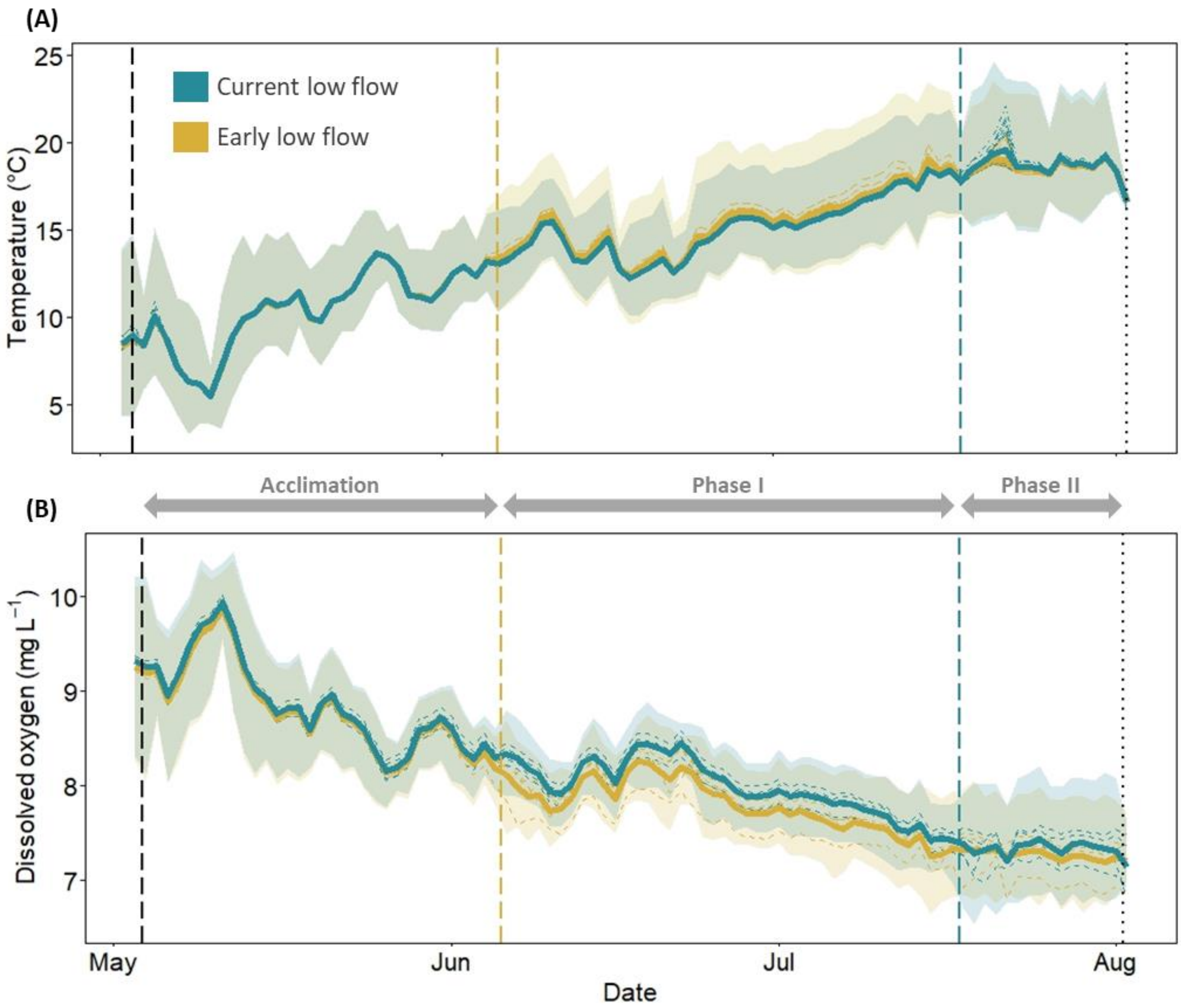

**Figure S2** Mean daily (solid lines) **(A)** temperature and **(B)** dissolved oxygen concentrations over the experiment. Channels exposed to *early* and *current* low-flow treatments are displayed in yellow and blue, respectively. Shaded area represents the daily range values (i.e., maximum and minimum). Dotted lines represent daily mean values for each experimental unit (temperature, n = 24) or channel (dissolved oxygen, n = 8). Vertical dotted lines represent the start and the end of the experiment (black), and the onset of low flows, colored by treatment.

**Table S1** Outputs from the repeated measure ANOVAs used to test the effects of low-flow duration (*early* vs. *current*) on temperature (ºC) and dissolved oxygen (DO, mg $L^{-1}$) over time. Analyses were performed separately for each phase of the experiment (*Acclimation*, *Phase I* and *Phase II*). Daily temperature mean and range (maximum - minimum) were calculated at the experimental unit level (n = 24 sensors, one per experiment unit), while DO mean and range were calculated at the channel level (n = 8 sensors, one per channel).

| **Response variable** | **Phase** | **Time (day)** | **Low-flow duration** | **Time × Low-flow** |
|---|---|---|---|---|
| Temperature (mean) | *Acclimation* (T1 – T4) | **$F_{6,776}$ = 39056, P < 0.001** | $F_{6,776}$ = 0.053, P = 0.825 | $F_{6,776}$ = 1.49, P = 0.268 |
| Temperature (range) | | **$F_{6,776}$ = 1203, P < 0.001** | $F_{6,776}$ = 0.20, P = 0.672 | $F_{6,776}$ = 1.48, P = 0.270 |
| Dissolved oxygen (mean) | | **$F_{6,248}$ = 2213, P < 0.001** | $F_{6,248}$ = 1.05, P = 0.345 | $F_{6,248}$ = 0.085, P = 0.780 |
| Dissolved oxygen (range) | | **$F_{6,248}$ = 15701, P < 0.001** | $F_{6,248}$ = 0.02, P = 0.895 | $F_{6,248}$ = 13.33, P = 0.011 |
| Temperature (mean) | *Phase I* (T5 – T10) | **$F_{6,992}$ = 6523, P < 0.001** | **$F_{6,992}$ = 18.19, P = 0.005** | $F_{6,992}$ = 0.01, P = 0.937 |
| Temperature (range) | | **$F_{6,992}$ = 58.46, P < 0.001** | **$F_{6,992}$ = 18.56, P = 0.005** | $F_{6,992}$ = 0.05, P = 0.826 |
| Dissolved oxygen (mean) | | **$F_{6,320}$ = 1254, P < 0.001** | $F_{6,320}$ = 3.83, P = 0.098 | $F_{6,320}$ = 2.19, P = 0.189 |
| Dissolved oxygen (range) | | $F_{6,320}$ = 4.95, P = 0.068 | **$F_{6,320}$ = 6.56, P = 0.043** | $F_{6,320}$ = 1.27, P = 0.303 |
| Temperature (mean) | *Phase II* (T11 – T12) | **$F_{6,368}$ = 11.50, P = 0.015** | $F_{6,368}$ = 1.59, P = 0.255 | $F_{6,368}$ = 0.87, P = 0.387 |
| Temperature (range) | | **$F_{6,368}$ = 53.45, P < 0.001** | $F_{6,368}$ = 3.51, P = 0.110 | $F_{6,368}$ = 0.45, P = 0.528 |
| Dissolved oxygen (mean) | | $F_{6,112}$ = 1.89, P = 0.218 | $F_{6,112}$ = 0.28, P = 0.616 | $F_{6,112}$ = 2.19, P = 0.189 |
| Dissolved oxygen (range) | | $F_{6,112}$ = 1.62, P = 0.336 | $F_{6,112}$ = 1.09, P = 0.336 | $F_{6,112}$ = 0.09, P = 0.775 |

**Table S2** Outputs from the linear (LM) and generalized linear models (GLM) used to test the effect of low-flow duration on fish growth and survival rates, respectively. The GLM was modeled as a binomial process with a logit link function. $R^2$ and pseudo-$R^2$ are displayed for LM and GLM, respectively. Significant results are shown in bold.

| **Response variable** | **Initial length** | **Low-flow duration** | **$R^2$** |
|---|---|---|---|
| Fish growth rate (SGR) | **$F_{1,17}$ = 12.37, P = 0.003** | $F_{1,17}$ = 2.72, P = 0.117 | 0.514 |
| Fish survival rate | | $F_{1,10}$ = 4.37, P = 0.063 | 0.183 |

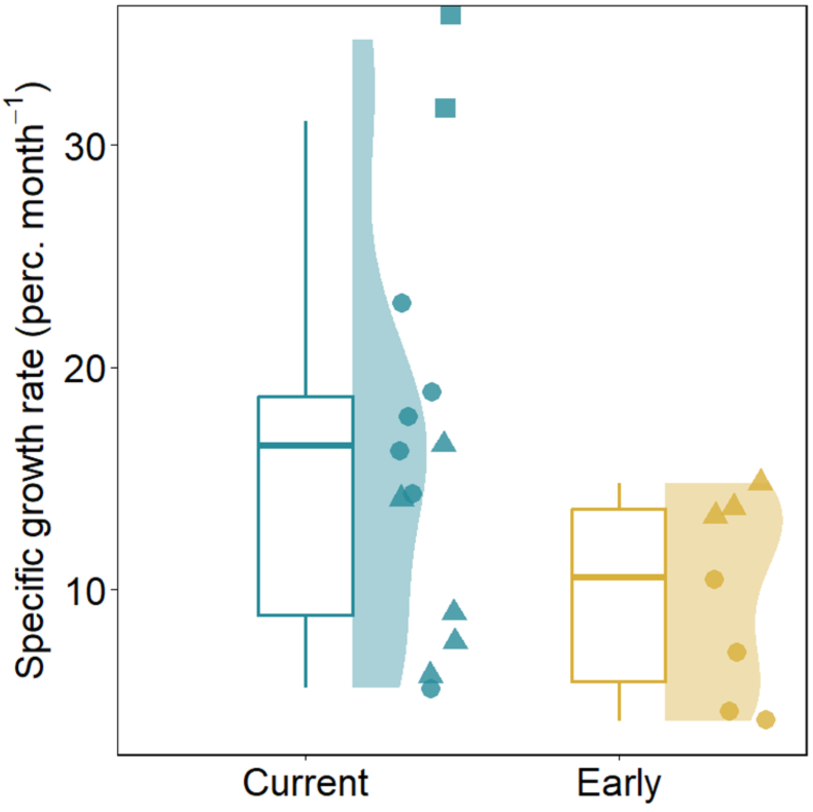

**Figure S3** Specific growth rate (% month$^{-1}$) of non-native Brown trout recaptured in *early* (yellow) and *current* (blue) low-flow conditions. Circles, triangles and squares represent large- ($n_{\text{Current low-flow}}$ = 6 and $n_{\text{Early low-flow}}$ = 4), medium- ($n_{\text{Current low-flow}}$ = 5 and $n_{\text{Early low-flow}}$ = 3) and small-sized ($n_{\text{Current low-flow}}$ = 2 and $n_{\text{Early-low flow}}$ = 0) individual trout, respectively. Boxplots and half violin plots represent the probability density of the data.

**Table S3** Outputs from the linear mixed effects models (LMMs) and PERMANOVAs used to test the effect of low-flow duration (*early* vs. *current*), presence of non-native trout (*fish* vs. *fishless*) and their interaction on emerging insect abundance, cumulative abundance (seasonally-aggregated), community structure (Bray-Curtis dissimilarity; PERMANOVA) and composition (Sørensen distance; PERMANOVA) and benthic algae biomass (including green algae, diatoms and cyanobacteria). For each LMM, conditional ($R^2C$, variance explained by fixed and random factors) and marginal ($R^2M$, variance explained by fixed factors) R-squared are displayed. Significant results are shown in bold.

| **Response variable** | **Phase (sampling)** | **Time (week)** | **Temperature** | **Introduced fish** | **Low-flow duration** | **Introduction × Low-flow** | **$R^2C$ / $R^2M$** |
|---|---|---|---|---|---|---|---|
| Abundance | | $F_{1,68.8}$ = 0.59, P = 0.445 | **$F_{1,68.8}$ = 4.50, P = 0.037** | $F_{1,71.4}$ = 0.39, P = 0.532 | $F_{1,5.5}$ = 0.26, P = 0.627 | $F_{1,70.5}$ = 0.25, P = 0.618 | 0.393 / 0.274 |
| Cum. abundance | *Acclimation* (T1 – T4) | $F_{1,68.1}$ = 3.02, P = 0.087 | $F_{1,68.2}$ = 3.19, P = 0.078 | $F_{1,71.6}$ = 0.95, P = 0.334 | $F_{1,5.9}$ = 0.33, P = 0.589 | $F_{1,71.8}$ = 0.58, P = 0.449 | 0.806 / 0.547 |
| Community structure | | — | F = 1.59, P = 0.190 | F = 2.21, P = 0.135 | F = 0.14, P = 0.944 | F = 1.37, P = 0.296 | — |
| Abundance | | $F_{1,121.7}$ = 0.70, P = 0.404 | **$F_{1,121.6}$ = 7.46, P = 0.007** | $F_{1,122.8}$ = 0.08, P = 0.775 | $F_{1,5.9}$ = 2.61, P = 0.159 | $F_{1,123.3}$ = 0.21, P = 0.649 | 0.326 / 0.282 |
| Cum. abundance | | **$F_{1,120.2}$ = 43.15, P < 0.001** | $F_{1,120.2}$ = 2.16, P = 0.145 | **$F_{1,122.4}$ = 19.00, P < 0.001** | $F_{1,6.0}$ = 1.99, P = 0.208 | $F_{1,122.3}$ = 3.05, P = 0.083 | 0.752 / 0.530 |
| Community structure | *Phase I* (T5 – T10) | — | **F = 10.15, P < 0.001** | F = 1.90, P = 0.095 | **F = 5.42, P = 0.004** | F = 1.86, P = 0.110 | — |
| Community composition | | — | **F = 3.37, P = 0.003** | F = 1.67, P = 0.107 | **F = 2.24, P = 0.026*** | F = 1.34, P = 0.217 | — |
| Abundance | | **$F_{1,36.2}$ = 41.62, P < 0.001** | $F_{1,39.5}$ = 0.07, P = 0.790 | $F_{1,36.6}$ = 3.97, *P = 0.054* | $F_{1,5.8}$ = 1.75, P = 0.236 | $F_{1,36.6}$ = 0.23, P = 0.637 | 0.787 / 0.325 |
| Cum. abundance | *Phase II* (T11 – T12) | **$F_{1,36.1}$ = 7.07 P = 0.012** | $F_{1,38.9}$ = 1.90, P = 0.176 | **$F_{1,36.5}$ = 8.94, P = 0.005** | $F_{1,5.9}$ = 0.01, P = 0.941 | $F_{1,36.5}$ = 0.01, P = 0.917 | 0.765 / 0.117 |
| Community structure | | — | **F = 2.67, P = 0.032** | F = 1.14, P = 0.301 | F = 1.61, P = 0.173 | F = 0.28, P = 0.957 | — |
| Community composition | | — | F = 1.29, P = 0.316 | F = 0.916, P = 0.462 | F = 1.88, P = 0.142 | F = 0.46, P = 0.776 | — |

| | | | | | | | |
|---|---|---|---|---|---|---|---|
| Algae biomass | (T12) | — | — | **$F_{1,43.8}$ = 24.13, P < 0.001** | $F_{1,5.6}$ = 0.91, P = 0.379 | $F_{1,43.8}$ = 0.29, P = 0.592 | _ / 0.368 |
| Green algae | | — | — | **$F_{1, 43.8}$ = 4.17, P = 0.047** | $F_{1, 5.6}$ = 0.01, P = 0.912 | $F_{1, 43.8}$ = 0.12, P = 0.731 | _ / 0.090 |
| Diatoms | | — | — | $F_{1, 43.6}$ = 1.62, P = 0.209 | $F_{1, 5.6}$ = 0.70, P = 0.437 | $F_{1, 43.6}$ = 0.51, P = 0.481 | _ / 0.061 |
| Cyanobacteria | | — | — | **$F_{1, 43.8}$ = 5.06, P = 0.030** | $F_{1, 5.6}$ = 0.24, P = 0.641 | $F_{1, 43.8}$ = 0.45, P = 0.504 | _ / 0.117 |

* Non-homogeneity of group dispersion (*betadisper*: P = 0.031)

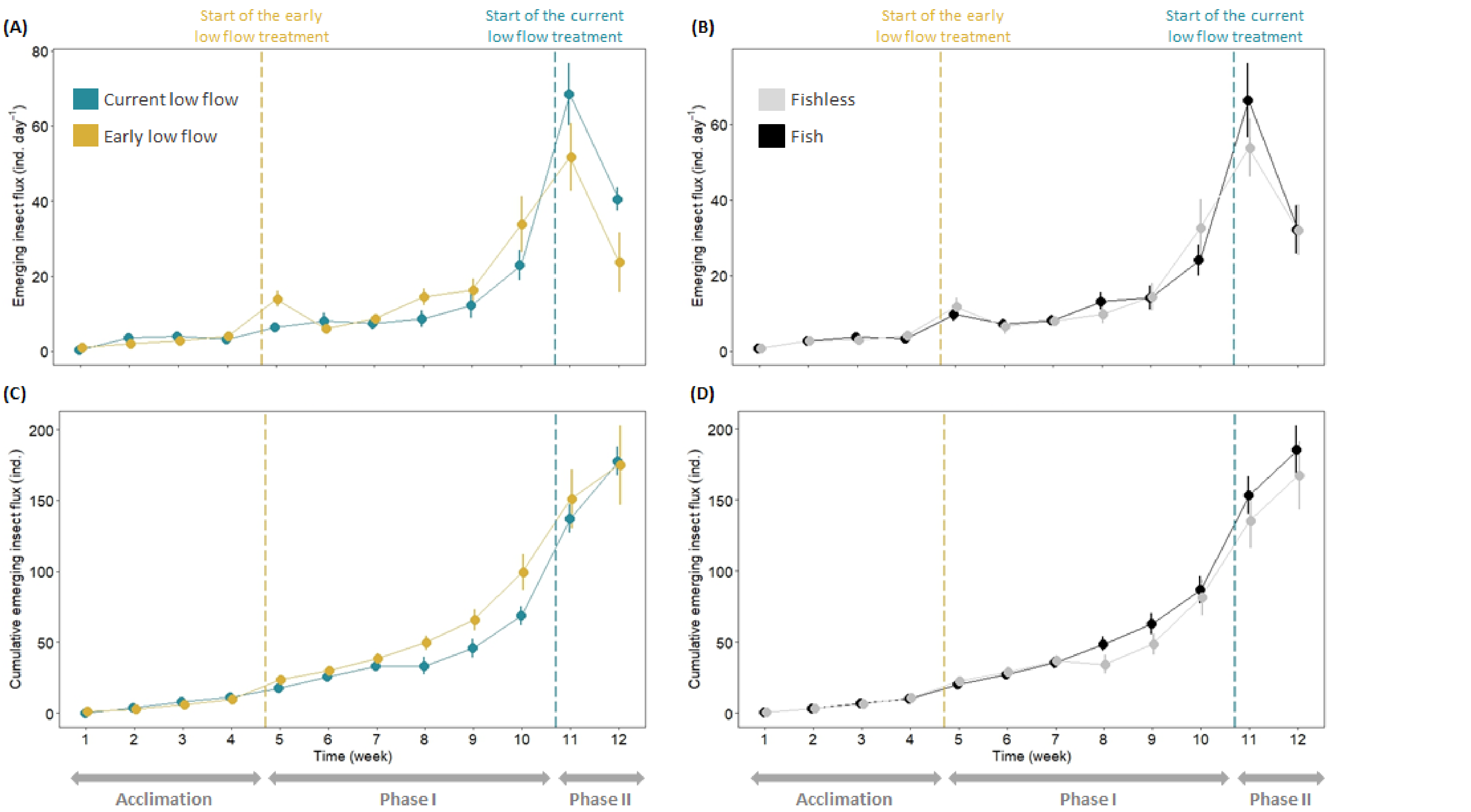

89 **Figure S4** Temporal variation in emerging insect abundances (mean ± SE) over the experiment and according to low-flow duration (left panels) and presence of non-native trout (right panel).

**Table S4** Outputs from the generalized linear mixed effects models (GLMMs) used to test the effect of low-flow duration (*early* vs. *current*), presence of non-native trout (*fish* vs. *fishless*) and their interaction on cumulative abundance (seasonally-aggregated) of the main Order, Family and feeding groups of emerging insects. Data from *Phase I* and *Phase II* were pooled together (T6, T9 and T12). GLMMs were fitted with negative binomial distributions, except the model with *Hydroptilidae* as a response variable which was fitted with a generalized Poisson distribution. Significant results are shown in bold.

| **Response variable** | **Time (week)** | **Temperature** | **Introduced fish** | **Low-flow duration** | **Introduction × Low-flow** |
|---|---|---|---|---|---|
| Diptera | $X^2$ = 0.54, P = 0.463 | **$X^2$ = 5.79, P = 0.016** | **$X^2$ = 7.78, P = 0.005** | **$X^2$ = 9.90, P = 0.002** | $X^2$ = 0.62, P = 0.435 |
| *Ceratopogonidae* | $X^2$ = 0.94, P = 0.332 | $X^2$ = 0.08, P = 0.773 | $X^2$ = 1.86, P = 0.173 | $X^2$ = 9.24, P = 0.337 | $X^2$ = 0.83, P = 0.362 |
| *Chironomidae* | $X^2$ = 0.52, P = 0.470 | **$X^2$ = 5.19, P = 0.023** | **$X^2$ = 10.14, P = 0.001** | **$X^2$ = 7.36, P = 0.007** | $X^2$ = 0.45, P = 0.502 |
| EPT | **$X^2$ = 4.26, P = 0.039** | $X^2$ = 0.31, P = 0.577 | **$X^2$ = 5.04, P = 0.025** | $X^2$ = 0.16, P = 0.689 | $X^2$ = 1.48, P = 0.223 |
| *Baetidae* | $X^2$ = 3.74, P = 0.053 | $X^2$ = 0.73, P = 0.392 | **$X^2$ = 7.16, P = 0.007** | $X^2$ = 0.01, P = 0.907 | $X^2$ = 3.41, P = 0.065 |
| *Leptophlebidae* | $X^2$ = 0.03, P = 0.852 | $X^2$ = 0.59, P = 0.442 | $X^2$ = 0.44, P = 0.507 | $X^2$ = 0.22, P = 0.639 | $X^2$ = 1.99, P = 0.158 |
| *Hydroptilidae* | $X^2$ = 0.16, P = 0.691 | $X^2$ = 0.39, P = 0.530 | **$X^2$ = 12.79, P < 0.001** | $X^2$ = 0.10, P = 0.748 | $X^2$ = 2.20, P = 0.138 |
| Scraper-grazers | $X^2$ = 0.01, P = 0.935 | $X^2$ = 0.45, P = 0.504 | **$X^2$ = 12.00, P = 0.001** | $X^2$ = 0.91, P = 0.340 | $X^2$ = 0.17, P = 0.677 |
| Predators | $X^2$ = 0.25, P = 0.619 | $X^2$ = 0.59, P = 0.443 | $X^2$ = 2.03, P = 0.154 | $X^2$ = 1.19, P = 0.275 | $X^2$ = 0.51, P = 0.475 |
| Collector-gatherers | $X^2$ = 0.84, P = 0.358 | $X^2$ = 3.75, P = 0.053 | **$X^2$ = 8.58, P = 0.003** | $X^2$ = 3.27, P = 0.070 | $X^2$ = 1.32, P = 0.250 |
| Shredders | $X^2$ = 0.78, P = 0.377 | $X^2$ = 0.55, P = 0.459 | $X^2$ = 0.16, P = 0.685 | **$X^2$ = 7.91, P = 0.005** | **$X^2$ = 6.72, P = 0.010** |

**Table S5** Outputs from the generalized linear mixed effects models (GLMMs) used to test for effects of low-flow duration (*early* vs. *current*), presence of non-native trout (*fish* vs. *fishless*), and their interaction, on abundance of emerging insects, grouping them in different ways. We ran tests at the Order and supra-Order level (i.e., Diptera and EPT - Ephemeroptera, Plecoptera and Trichoptera), at the Family level (i.e., Chironomidae, Ceratopogonidae, Baetidae, Leptophlebiidae, and Hydroptilidae), and at the functional feeding group level (i.e., Scraper-grazers, Predators, Collector-gatherers, and Shredders). Data from *Phase I* and *Phase II* were pooled together (T6, T9 and T12). GLMMs were fitted with negative binomial distributions. Significant results are shown in bold.

| **Response variable** | **Time (week)** | **Temperature** | **Introduced fish** | **Low-flow duration** | **Introduction × Low-flow** |
|---|---|---|---|---|---|
| Diptera | $X^2 = 0.36$, P = 0.548 | **$X^2 = 8.69$, P = 0.003** | **$X^2 = 4.45$, P = 0.035** | $X^2 = 1.90$, P = 0.168 | $X^2 = 0.60$, P = 0.440 |
| *Ceratopogonidae* | $X^2 = 0.28$, P = 0.599 | $X^2 = 0.18$, P = 0.672 | $X^2 = 1.16$, P = 0.280 | $X^2 = 0.23$, P = 0.630 | $X^2 = 0.21$, P = 0.647 |
| *Chironomidae* | $X^2 = 0.11$, P = 0.745 | **$X^2 = 6.86$, P = 0.009** | **$X^2 = 6.09$, P = 0.014** | $X^2 = 1.68$, P = 0.195 | $X^2 = 0.60$, P = 0.437 |
| EPT | $X^2 = 0.87$, P = 0.351 | $X^2 = 0.09$, P = 0.764 | $X^2 = 0.99$, P = 0.320 | $X^2 = 0.02$, P = 0.889 | $X^2 = 0.27$, P = 0.604 |
| *Baetidae* | $X^2 = 2.39$, P = 0.122 | $X^2 = 1.06$, P = 0.303 | $X^2 = 2.05$, P = 0.152 | $X^2 = 0.03$, P = 0.861 | $X^2 = 0.81$, P = 0.369 |
| *Leptophlebiidae* | $X^2 = 2.88$, P = 0.090 | $X^2 = 3.35$, P = 0.067 | $X^2 = 0.46$, P = 0.490 | $X^2 = 0.18$, P = 0.668 | $X^2 = 0.91$, P = 0.339 |
| *Hydroptilidae* | $X^2 = 0.02$, P = 0.900 | $X^2 = 0.49$, P = 0.483 | **$X^2 = 4.78$, P = 0.029** | $X^2 = 0.04$, P = 0.839 | $X^2 = 0.62$, P = 0.432 |
| Scraper-grazers | $X^2 = 0.86$, P = 0.354 | $X^2 = 1.43$, P = 0.231 | $X^2 = 2.55$, P = 0.110 | $X^2 = 0.20$, P = 0.654 | $X^2 = 0.11$, P = 0.744 |
| Predators | $X^2 = 0.04$, P = 0.843 | $X^2 = 0.59$, P = 0.441 | $X^2 = 1.33$, P = 0.249 | $X^2 = 0.45$, P = 0.503 | $X^2 = 0.07$, P = 0.787 |
| Collector-gatherers | $X^2 = 0.01$, P = 0.941 | **$X^2 = 4.83$, P = 0.028** | **$X^2 = 4.42$, P = 0.036** | $X^2 = 0.75$, P = 0.385 | $X^2 = 1.38$, P = 0.240 |
| Shredders | $X^2 = 0.29$, P = 0.593 | $X^2 = 3.02$, P = 0.023 | $X^2 = 0.06$, P = 0.800 | $X^2 = 3.10$, P = 0.078 | **$X^2 = 4.76$, P = 0.029** |

**Table S6** Results of Tukey post-hoc analyses used to test the effects of significant interactions between low-flow duration (*current* vs. *early*) and presence of non-native trout fish (*fish* vs. *fishless*) on the cumulative abundance and abundance of shredders (results reported in Table S4 and S5). Significant results are shown in bold.

| **Cumulative abundance of Shredders** | | | |
|---|---|---|---|
| **Comparisons** | **Estimate (SE)** | **z-value** | **P-value** |
| Current-Fishless vs. Current-Fish | 0.88 (0.30) | -0.39 | 0.698 |
| Early-Fishless vs. Early-Fish | 2.96 (0.98) | 3.27 | **0.001** |
| Current-Fish vs. Early-Fish | 0.37 (0.13) | -2.76 | **0.006** |
| Current-Fishless vs. Early-Fishless | 1.24 (0.48) | 0.55 | 0.583 |
| **Abundance of Shredders** | | | |
| **Comparisons** | **Estimate (SE)** | **z-value** | **P-value** |
| Current-Fishless vs. Current-Fish | 0.91 (0.34) | -0.25 | 0.800 |
| Early-Fishless vs. Early-Fish | 2.91 (1.09) | 2.85 | **0.004** |
| Current-Fish vs. Early-Fish | 0.51 (0.20) | -1.76 | 0.078 |
| Current-Fishless vs. Early-Fishless | 1.62 (0.65) | 1.21 | 0.227 |

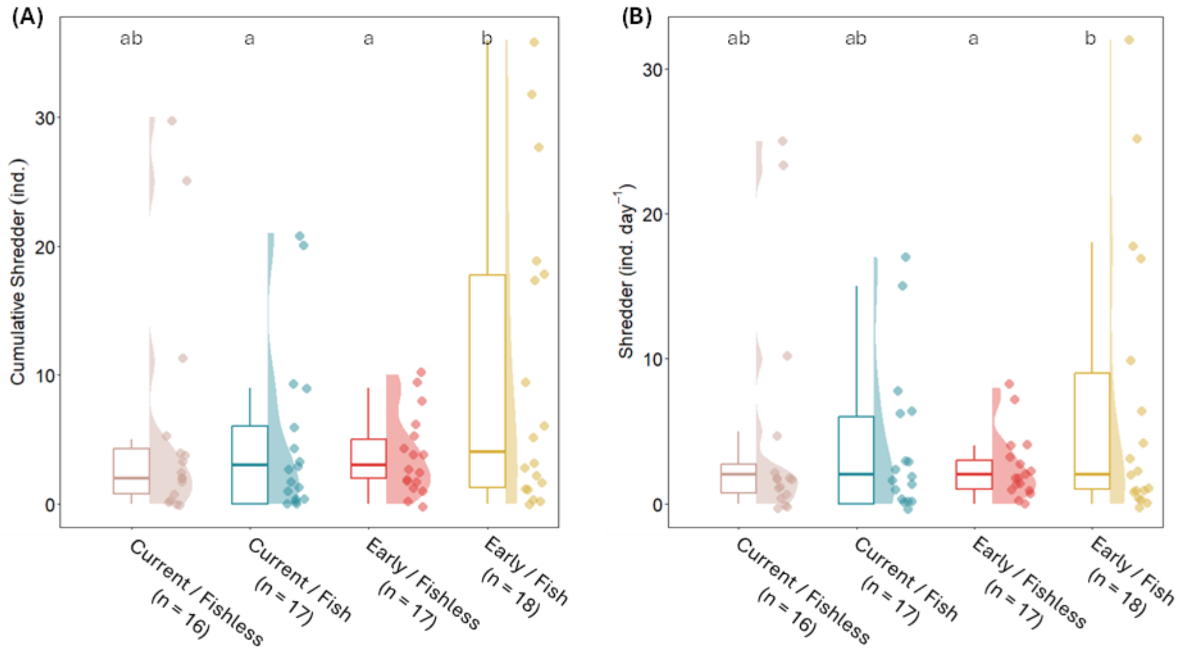

**Figure S5** Raincloud plots showing the effect of treatment combinations on (A) the cumulative abundance and (B) the abundance of shredders collected during *Phase I* and *Phase II*. Dots represent the samples , boxplots and half violin plots represent the probability density of the data. Letters indicate significant differences.

**Table S7** Effects of low-flow duration on the abundance of emerging insects that significantly explained Bray-Curtis dissimilarity between *early* and *current* low-flow conditions during the *Phase I* of the experiment (samples collected at T6 and T9). Results are from GLMMs fitted with negative binomial distributions and with stream identity as a random effect. Significant results ($P < 0.05$) are shown in bold. Asterisks indicate P = 0.001 (***) and $P < 0.05$ (*) for analyses associated with community dissimilarity.

| **Taxa** | **Low-flow duration** | **Mean ± SD in *Current* vs. *Early* low flows** |
|---|---|---|
| *Orthocladiinae**** | **$X^2$ = 5.24, P = 0.022** | 2.67 ± 4.78 vs. 6.44 ± 6.32 |
| *Chironominae**** | $X^2$ = 0.05, P = 0.827 | 1.14 ± 1.71 vs. 1.26 ± 2.00 |
| *Prodiamesinae**** | **$X^2$ = 12.50, P < 0.001** | 0.24 ± 0.89 vs. 1.78 ± 2.15 |
| *Diphetor hageni** | $X^2$ < 0.01, P = 0.968 | 0.57 ± 0.98 vs. 0.26 ± 0.54 |
| *Baetis** | $X^2$ = 1.74, P = 0.187 | 0.38 ± 0.80 vs. 0.39 ± 0.78 |
| *Lepidostomatidae** | $X^2$ = 0.48, P = 0.487 | 0.10 ± 0.30 vs. 0.17 ± 0.39 |

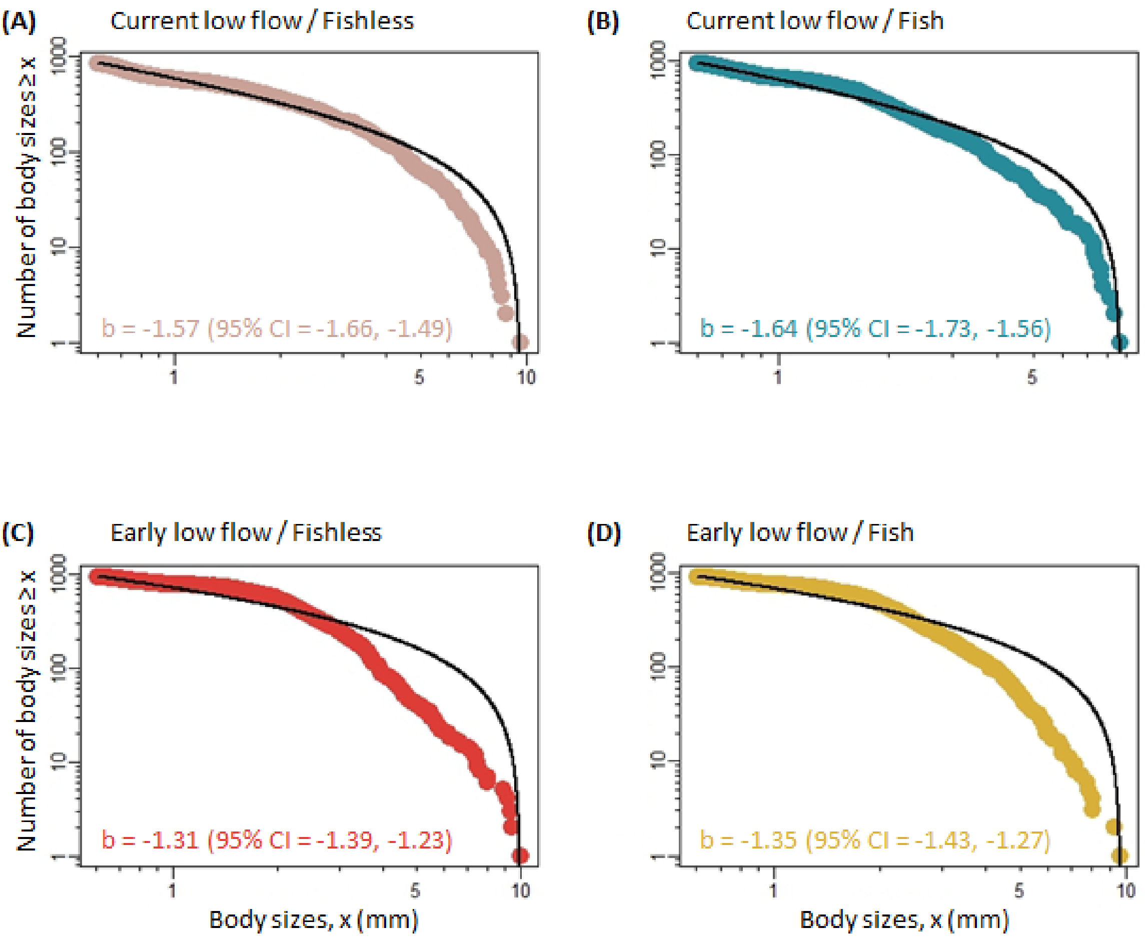

**Figure S6** Relationships between body size (x, mm) and the number of insects with body size ≥ x on logarithmic scales. The rank frequency plots visualise the fit of the size spectra slopes using maximum likelihood estimation of a bounded power-law distribution for each treatment combination. Size spectra slopes (b) and 95% CI are provided at the bottom of each panel.
